# iPSC-derived astrocytes to model phenotype-specific differential neuroinflammatory and metabolic responses in X-linked adrenoleukodystrophy

**DOI:** 10.1101/2022.09.09.507263

**Authors:** Parveen Parasar, Navtej Kaur, Laila M Poisson, Jaspreet Singh

## Abstract

X-linked adrenoleukodystrophy (X-ALD) is an inherited progressive metabolic disorder caused by pathogenic variants in the *ABCD1* gene, which leads to accumulation of very long chain fatty acids in body fluids and tissues including brain and spinal cord. In the absence of a clear genotype-phenotype correlation the molecular mechanisms of the severe cerebral adrenoleukodystrophy (cALD) and the milder adrenomyeloneuropathy (AMN) phenotypes remain unknown. Given our previous evidence of role of astrocytes in the neuroinflammatory response in X-ALD we investigated the metabolic and molecular profiles of astrocytes derived from induced pluripotent stem cells (iPSC). The iPSCs were in turn generated from skin fibroblasts from healthy controls and patients with AMN or cALD. AMN and cALD astrocytes exhibited lack of *ABCD1* and accumulation of very long chain fatty acids, a hallmark of X-ALD disease. Further, cALD astrocytes harbor significantly higher phosphorylation of STAT3, increased Toll-like receptor expression and higher chemokine and cytokine expression. In this first report of miRNA sequencing in X-ALD astrocytes, we observed that miR-9 expression was associated with increasing disease severity phenotype. CRISPR-Cas9 knock-in of *ABCD1ABCD1* gene expression differentially affected the expression of key molecular, metabolic and microRNA targets in AMN and cALD astrocytes. Extensive characterization of the AMN and cALD iPSC-derived astrocyte model demonstrates critical aspects of X-ALD inflammatory disease in response to *ABCD1ABCD1* mutation and can be further utilized for exploring the contribution of astrocytes to differential inflammatory response in cALD.

## INTRODUCTION

X-linked adrenoleukodystrophy (X-ALD) is a progressive peroxisomal disease due to a defective *ABCD1* gene encoding the peroxisomal ABC half-transporter *ABCD1* or adrenoleukodystrophy (ALD) protein. Deleterious and/or pathogenic variants of *ABCD1* impair β-oxidation of fatty acids, leading to accumulation of very long chain fatty acids (VLCFAs), predominantly C26:0 in blood and tissues, which adversely affects the brain, spinal cord, and adrenals.(Moser, 1995) A total of 1000 unique variants of *ABCD1ABCD1* in >3400 cases *ABCD1ABCD1* variants have been reported in X-ALD; however, no correlation of genotype to clinical phenotype exists (https://adrenoleukodystrophy.info/) (Kemp et al., 2001; Mallack et al., 2022).

X-ALD phenotypes include rapidly progressive inflammatory demyelinating cerebral adrenoleukodystrophy (cALD), milder adult-onset forms, adrenomyeloneuropathy (AMN), and adrenal insufficiency without neurologic involvement. Approximately 35% of affected males develop cALD before reaching 12 years of age. Without early intervention, most patients with cALD die within a decade after diagnosis. AMN is a gradually developing phenotype, first affecting males with dysfunctional *ABCD1* at age 20-30 years. AMN may develop as the pure non-inflammatory axonopathy form, manifesting with gait disturbances and bladder dysfunction, or as the AMN-cerebral form, manifesting with cerebral inflammation in addition to the axonopathy AMN form. About 30% of patients with AMN develop cerebral inflammation, ultimately progressing to the fatal cALD form in adulthood (Kemp *et al*., 2001; Moser, 1995).

Human astrocytes are the major cell population in the central nervous system and play a substantial role in X-ALD pathogenesis, and the peroxisomal *ABCD1* protein plays a role in the murine astrocyte inflammatory response (Singh et al., 2009) We previously documented that loss of adenosine monophosphate-activated protein kinase α1 (AMPKα1) in patients with cALD was associated with a higher inflammatory profile in patient-derived cells (Singh and Giri, 2014; Singh et al., 2016; Singh et al., 2015); however, the underlying definitive mechanisms of phenotypic variability and differential inflammatory responses between AMN and cALD phenotypes remain unknown.

Current mouse models (Forss-Petter et al., 1997; Kobayashi et al., 1997; Lu et al., 1997) poorly represent clinical X-ALD phenotypic heterogeneity, and mice do not develop neurologic phenotypes representative of human cALD phenotypes. This begs a clear need for a directly translatable cellular model to study X-ALD disease mechanisms. In the context of limited models and inaccessibility of human astrocytes, using induced pluripotent stem cells (iPSCs) from patients is an emerging paradigm to study etiopathogenetic mechanisms at the cellular level *in vitro*, screen potential therapeutics, and provide defined conditions for reproducibly performing research (Marchetto et al., 2011). Furthermore, iPSCs retain pluripotency, the ability to differentiate into any cell type, and they carry the genetic background of patients, including disease-causing genetic variants (Park et al., 2008).

Here we found that reprogrammed iPSCs from healthy control (CTL), AMN, and cALD fibroblasts can be differentiated into astrocytes and document many hallmarks of cALD and AMN phenotype. We document novel insights into differential inflammatory, metabolic and microRNA response in human iPSC-derived astrocytes from patients with cALD and AMN phenotypes. These iPSC-derived astrocytes provide a valuable model to investigate differential disease mechanism in AMN and cALD. Furthermore, Lentiviral delivery and knock-in of *ABCD1ABCD1* gene in AMN and cALD astrocytes led to differential reversal of mitochondrial, glycolytic, and inflammatory profile in cALD and AMN astrocytes highlighting the role of pathways beyond *ABCD1* in the differential inflammatory response.

## RESULTS

### Generation and Characterization of iPSCs and Differentiated Astrocytes

iPSCs were generated from CTL, AMN and cALD fibroblasts and expressed pluripotency markers OCT4, Nanog, Sox2, and TRA-1-60 (Figure 1A). Astrocytes differentiated from iPSCs expressed multiple neuronal cell markers such as glial fibrillary acidic protein (GFAP), nestin, aquaporin 4, TUJ1, S100 calcium-binding protein β (S100-β), aldehyde dehydrogenase family 1 member L1 (ALDH1L1), EAAT1, and A2B5(Figure 1B). ALDH1L1, S100-β, and aquaporin4 are markers of mature astrocytes. Expression of GFAP, S100-β, EAAT1, AQP4, and ALDH1markers was confirmed by immunoblot and qRT-PCR (Figure 1C-D). The level of S100-β protein was decreased in cALD and AMN astrocytes compared to CTL. AMN and cALD astrocytes showed higher AQP4 expression (Figure 1C).

**Figure 1:**
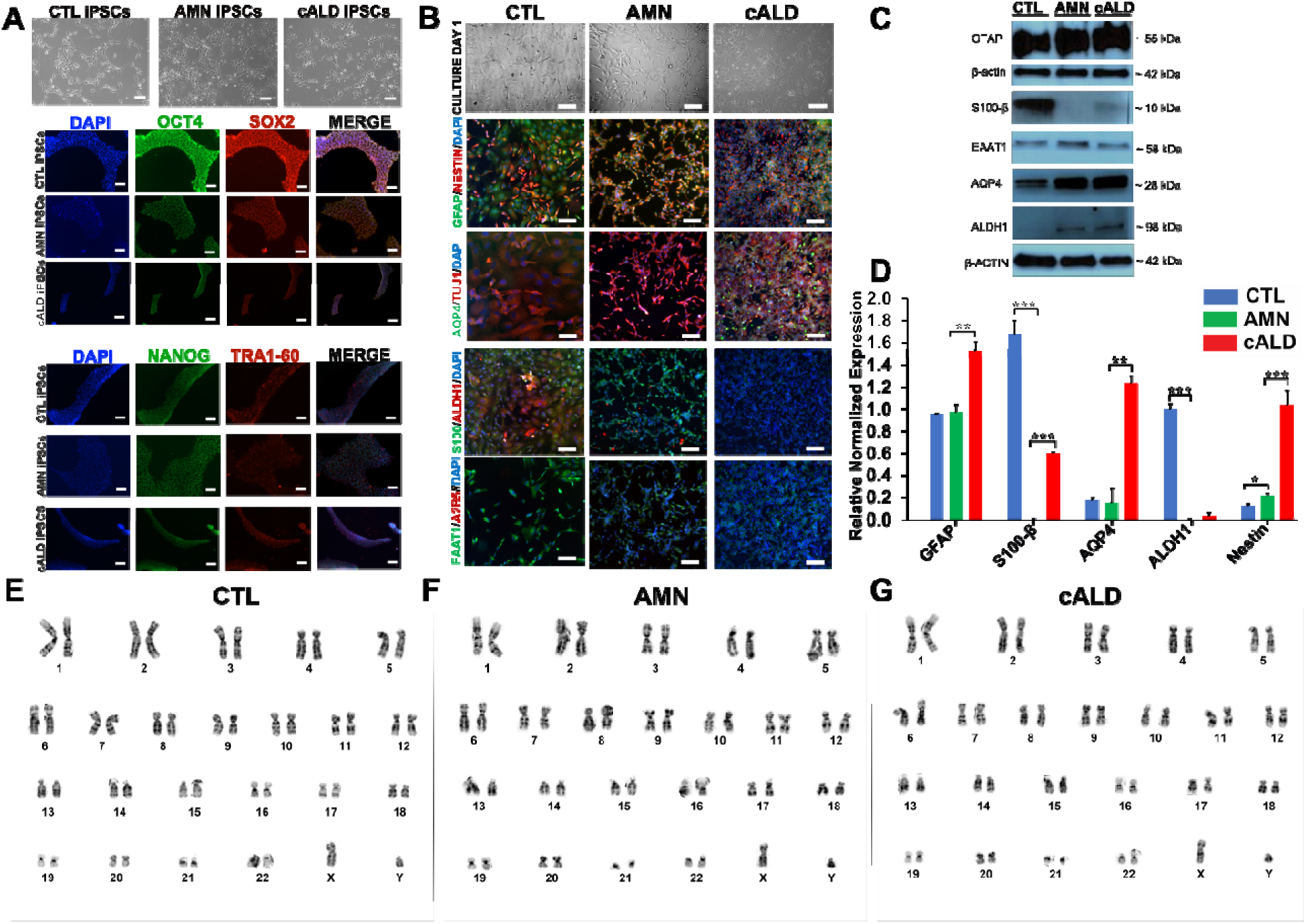
X-ALD fibroblasts can be successfully reprogrammed into iPSCs and thence differentiated into patient astrocytes. (A) Representative phase contrast (upper panel) and immunostained images of CTL, AMN, and cALD iPSCs. Upper panel shows morphology in culture (day 3) of reprogrammed CTL, AMN, and cALD iPSCs. Middle and lower panel represent phase contrast images for immunostaining for pluripotency markers: CTL, AMN, and cALD iPSCs with positive staining of OCT4 (green), SOX2 (red), NANOG (green), and TRA1-60 (red)(scale bars 150 µm). Nuclei are stained with DAPI (blue). (B) Phase contrast images (upper panel) of differentiated CTL, AMN, and cALD astrocytes in culture (day 1) and immunofluorescence staining (lower panels) with astrocyte specific markers: CTL, AMN, and cALD astrocytes stained positive with GFAP/Nestin (green/red), S100-β/ALDH1 (green/red), AQP4/tubulin βIII (green/red), and EAAT1/A2B5 (green/red)(scale bars 150 µm). Nuclei are stained with DAPI (blue). (C) Representative images for Western blotting with GFAP, S100-β, EAAT1, AQP4, ALDH1, and β-actin in CTL, AMN, and cALD patient astrocytes. (D) RT-qPCR quantification of gene expression for GFAP, S100-β, AQP4, ALDH1, and Nestin for astrocytes differentiated from CTL, AMN, and cALD iPSCs-derived astrocytes (n = 3). The gene expression of the individual sample was assessed with fold change using the comparative ΔΔ*Ct* method by normalizing with reference gene L-27 in CTL, AMN, and cALD astrocytes. Results represent means ± standard error from triplicates of each sample type. Two-tailed Student’s *t* test was used for data presented in 1D. Significance was assigned denoted for P-values of less than 0.05. **P < 0.01; ***P < 0.001. (E-G) Karyotype analysis for differentiated astrocytes from reprogrammed CTL (E), AMN (F), and cALD (G) fibroblasts. **Abbreviations**: AMN: Adrenomyeloneuropathy; cALD: Cerebral adrenoleukodystrophy; CTL: Control; iPSC: Induced pluripotent stem cells.

### Karyotype Analyses

CTL, AMN, and cALD astrocytes displayed normal male karyotypes and G-banding patterns with no structural or numerical changes, suggesting that reprogramming and differentiation did not induce chromosomal abnormalities in AMN and cALD astrocytes (Figure 1E-G). No anomalies or copy number changes were seen in any astrocyte groups per chromosome microarray (data not shown).

### AMN and cALD Astrocytes Show Loss of *ABCD1* Protein

We next investigated the status of *ABCD1* in CTL, AMN and cALD astrocytes. There was complete loss of *ABCD1 (ALDP) protein* in AMN and cALD astrocytes relative to CTL astrocytes. However, AMN and cALD astrocytes had a marked compensatory increase in *ABCD2* protein expression. ABCD3 was expressed at low levels and was not significantly different between CTL, AMN and cALD (Figure 2A).

**Figure 2:**
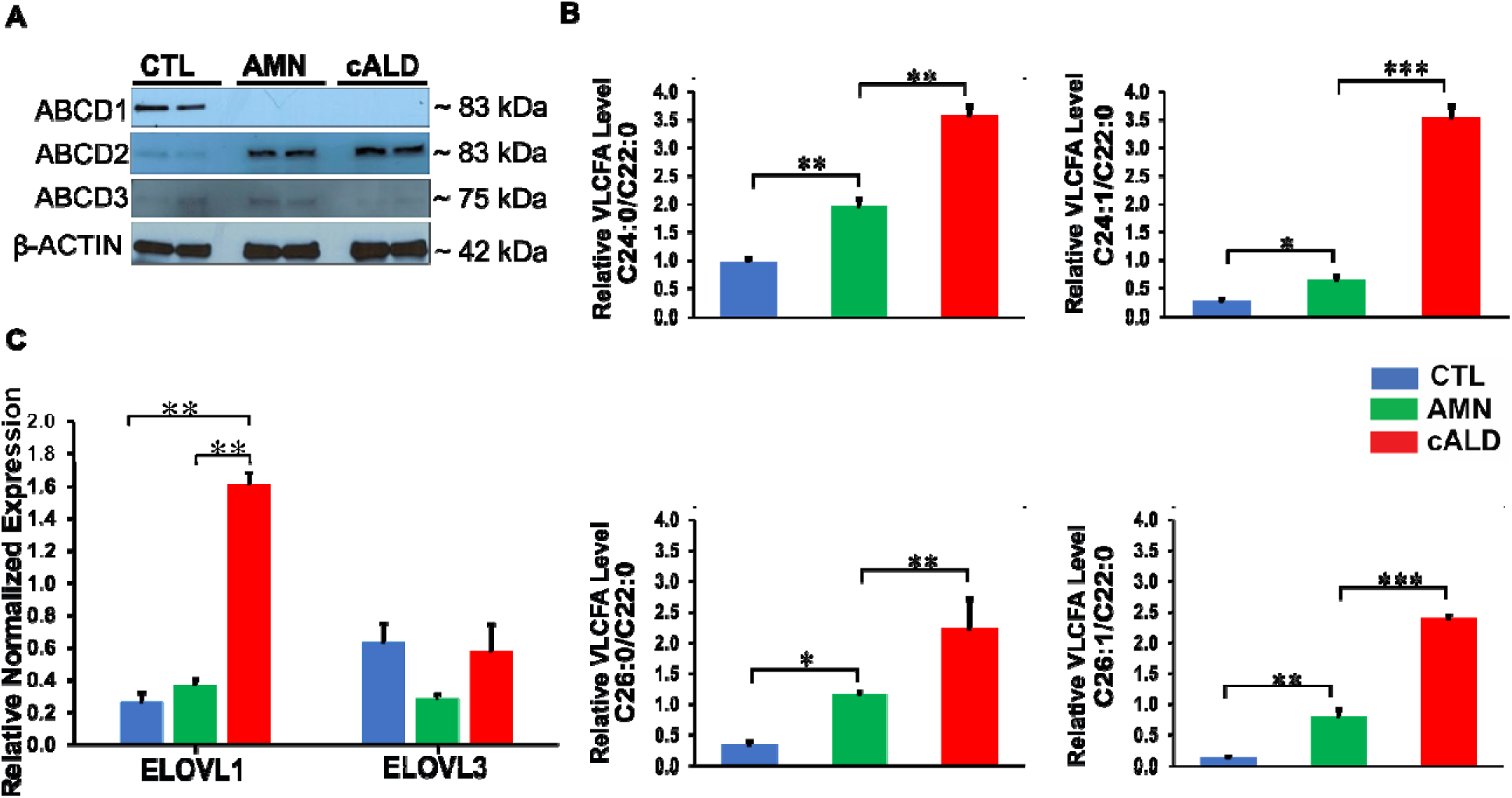
CTL, AMN, and cALD patient astrocytes retain biochemical characteristics of X-ALD patients. (A) Representative images for Western blotting with *ABCD1*, *ABCD2*, and *ABCD3*, and β-actin in CTL, AMN, and cALD patient astrocytes. (B) Quantification of VLCFA (ratio C24:0/C22:0 and C24:1/C22:0; C26:0/C22:0 and C26:1/C22:0) in astrocytes generated from CTL, AMN, and cALD patient fibroblast-derived iPSC-astrocytes. (C) RT-qPCR quantification of gene expression for ELOVL1 and ELOVL3 for astrocytes differentiated from CTL, AMN, and cALD iPSC-derived astrocytes (n = 3 technical replicates per line for astrocytes). The gene expression of the individual sample was assessed with fold change using the comparative ΔΔ*Ct* method by normalizing with reference gene L-27 in CTL, AMN, and cALD astrocytes. Results (Figure B and C) represent means ± standard error from triplicates of each of sample type. Two-tailed Student’s *t* test was used for data analysis. Significance was assigned for P-values of less than 0.05. **P < 0.01; ***P < 0.001. **Abbreviations**: AMN: Adrenomyeloneuropathy; cALD: Cerebral adrenoleukodystrophy; CTL: Control; VLCFA: Very long chain fatty acids.

### AMN and cALD Astrocytes Accumulated Higher VLCFAs

Peroxisomal VLCFA accumulation is determined by ALDP expression. Accumulation of VLCFA, the biochemical hallmark of X-ALD, was confirmed in AMN and cALD astrocytes. AMN astrocytes displayed significantly higher levels of saturated (C24:0, C26:0) and monounsaturated (C24:1, C26:1) (C24:0 and C26:1 (p<0.01); C26:0 and C24:1 (p<0.05)) VLCFA than CTL astrocytes, indicating impaired VLCFA β-oxidation (Figure 2B). cALD astrocytes showed further significantly increased saturated (p<0.01) and monounsaturated (p<0.001) VLCFA levels than AMN astrocytes (Figure 2B).

ELOVL1 catalyzes synthesis of saturated and monounsaturated VLCFAs (Ohno et al., 2010). cALD astrocytes showed approximately four-fold higher ELOVL1 (p < 0.01) expression than AMN and CTL astrocytes, and expression between AMN and CTL astrocytes did not differ. Expression of ELOVL3 did not differ significantly between the three astrocyte groups (Figure 2C).

### cALD Astrocytes Show Increased Basal Respiration and Elevated ATP-Linked OCR

VLCFA accumulation causes metabolic alterations in X-ALD (Singh and Giri, 2014; Singh *et al*., 2016; Singh *et al*., 2015). We performed mitochondrial extracellular flux analysis to assess OCR and ECAR in iPSC-derived astrocytes. The Cell Mito Stress Test showed that basal OCR (Figure 3A), non-mitochondrial OCR (Figure 3B), ATP-linked OCR (Figure 3C), maximal respiration, and spare capacity (Figure 3D) (p < 0.001) were higher in cALD astrocytes than in AMN astrocytes, indicating higher uncoupling and energy needs. Seahorse ATP rate assay showed that both AMN and cALD astrocytes had significantly lower ATP production (p < 0.05) than CTL astrocytes, but cALD astrocytes had a significantly higher ATP production rate (p < 0.05) than AMN astrocytes (Figure 3E).

**Figure 3:**
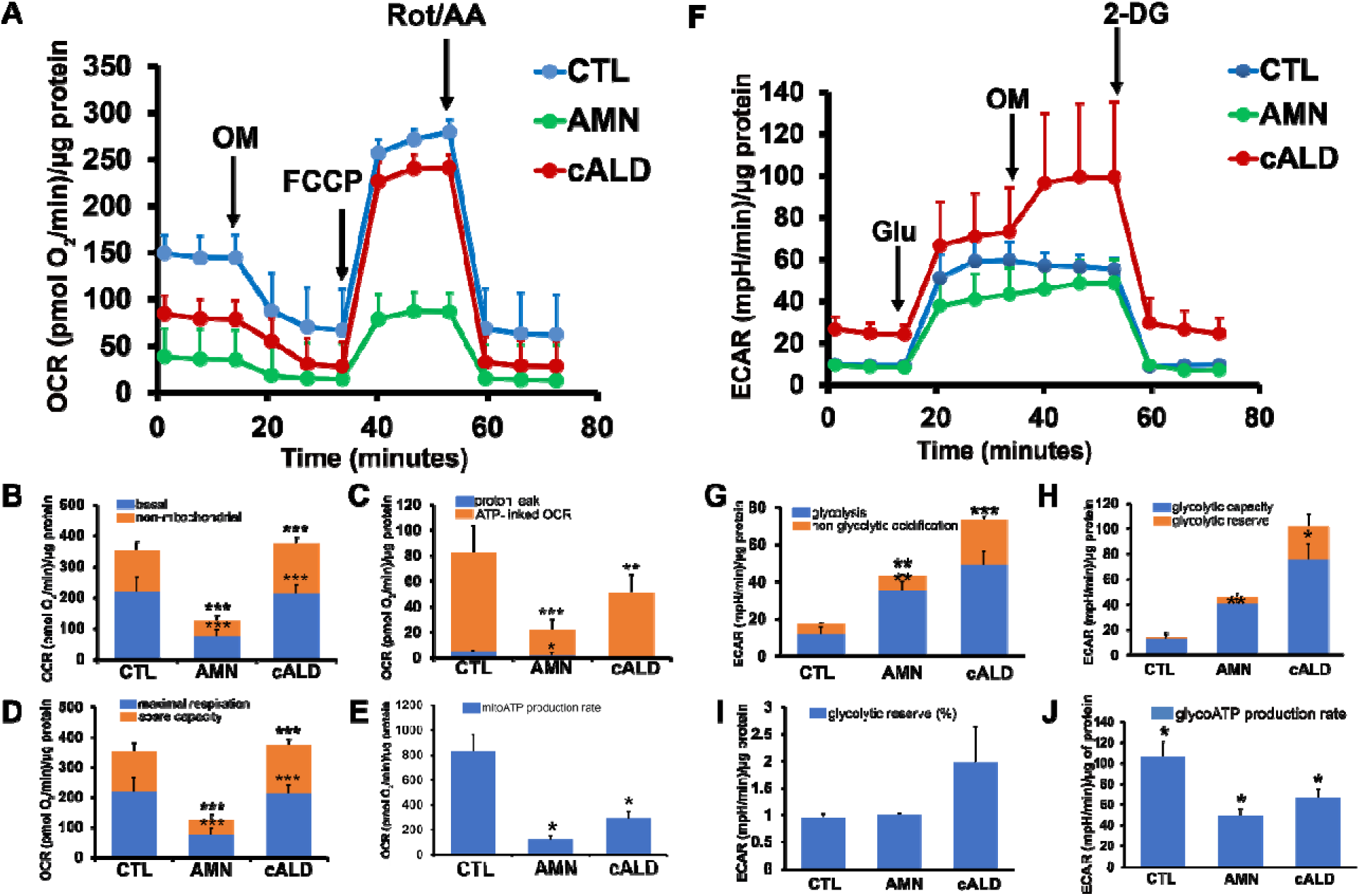
cALD astrocytes show altered parameters of OCR and ECAR in cell mito stress and glycolytic stress test assays. (A) Plot showing overall OCR in CTL, AMN, and cALD astrocytes. (B) Quantification of basal and non-mitochondrial OCR. (C) Quantification of proton leak and ATP-linked OCR. (D) Quantification of maximal respiration and spare respiratory capacity. (E) Quantification of mitochondrial ATP production rate (pmol ATP/minute). (F) Plot showing the overall ECAR in CTL, AMN, and cALD astrocytes. (G) Quantification of glycolysis and non-glycolytic acidification. (H) Quantification of glycolytic capacity and glycolytic reserve. (I) Quantification of percent glycolytic reserve. (J) Quantification of glycolytic ATP production rate (pmol ATP/minute). **Data information**: Results represent means ± standard error (n = 3 technical replicates) of each of sample type. Analysis was performed using the Wave software (version 2.6.1.53; Agilent) and normalized to the cell protein content (A-D and F-I). Data analysis in Figure E and J was performed after acquisition of data using XF Real-Time ATP rate assay generator (Agilent)(n = 3 technical replicates) for each sample type. Unpaired two-tailed Student’s *t* test or Mann-Whitney test were used for analysis. Significance was denoted for P-values of less than 0.05. **P < 0.01; ***P < 0.001. Significance level over AMN bars represent difference between CTL and AMN astrocyte samples and that over cALD bars represent the difference between AMN and cALD astrocyte samples. **Abbreviations**: AMN: Adrenomyeloneuropathy; cALD: Cerebral adrenoleukodystrophy; CTL: Control; ECAR: Extracellular acidification rate; OCR: Oxygen consumption rate; OM: Oligomycin; FCCP: Carbonyl cyanide-4 (trifluoromethoxy) phenylhydrazone; Rot/AA: Rotenone and antimycin-A; Glu: Glucose; OM: Oligomycin; 2-DG: 2-deoxi-D-glucose.

Intriguingly, we observed elevated ECAR in cALD astrocytes (Figure 3F), suggesting enhanced glycolytic activity. Glycolysis stress test revealed higher non-glycolytic ECAR (p < 0.001) and glycolysis (Figure 3G) as well as significantly higher glycolytic capacity (p < 0.05) and glycolytic reserve (p < 0.05) in cALD astrocytes (Figure 3H-I) than in AMN and CTL astrocytes, suggesting increased uncoupling in cALD cells. The higher energy demands of cALD astrocytes may indicate a proinflammatory state, showing that they are metabolically plastic. Lastly, while cALD astrocytes had a significantly lower glycolytic ATP (p < 0.05) production rate than AMN astrocytes, both AMN and cALD astrocytes had significantly lower glycolytic ATP production (p < 0.05) rates than CTL astrocytes (Figure 3J).

### Electron Microscopy Reveals Mitochondrial Structure Differences

To determine if mitochondrial functional changes were also associated with structural alterations, we performed electron microscopy to assess differences in cellular and mitochondrial structure between the CTL, AMN and cALD astrocytes. Importantly, AMN and cALD astrocytes contained more fatty acid droplets than CTL astrocytes (Figure 4A1-A3). We also observed disrupted cristae structure in cALD astrocytes relative to CTL and AMN astrocytes (Figure 4A4-A9). Dynamin related protein-1 (DRP-1), a key mitochondrial fission regulator, was elevated (p<0.01) in both AMN and cALD astrocytes, indicating mitochondrial fragmentation (Figure 4B-C). These findings suggest that cALD astrocytes have altered mitochondrial structure that may contribute to astrocytic mitochondrial dysfunction in X-ALD.

**Figure 4:**
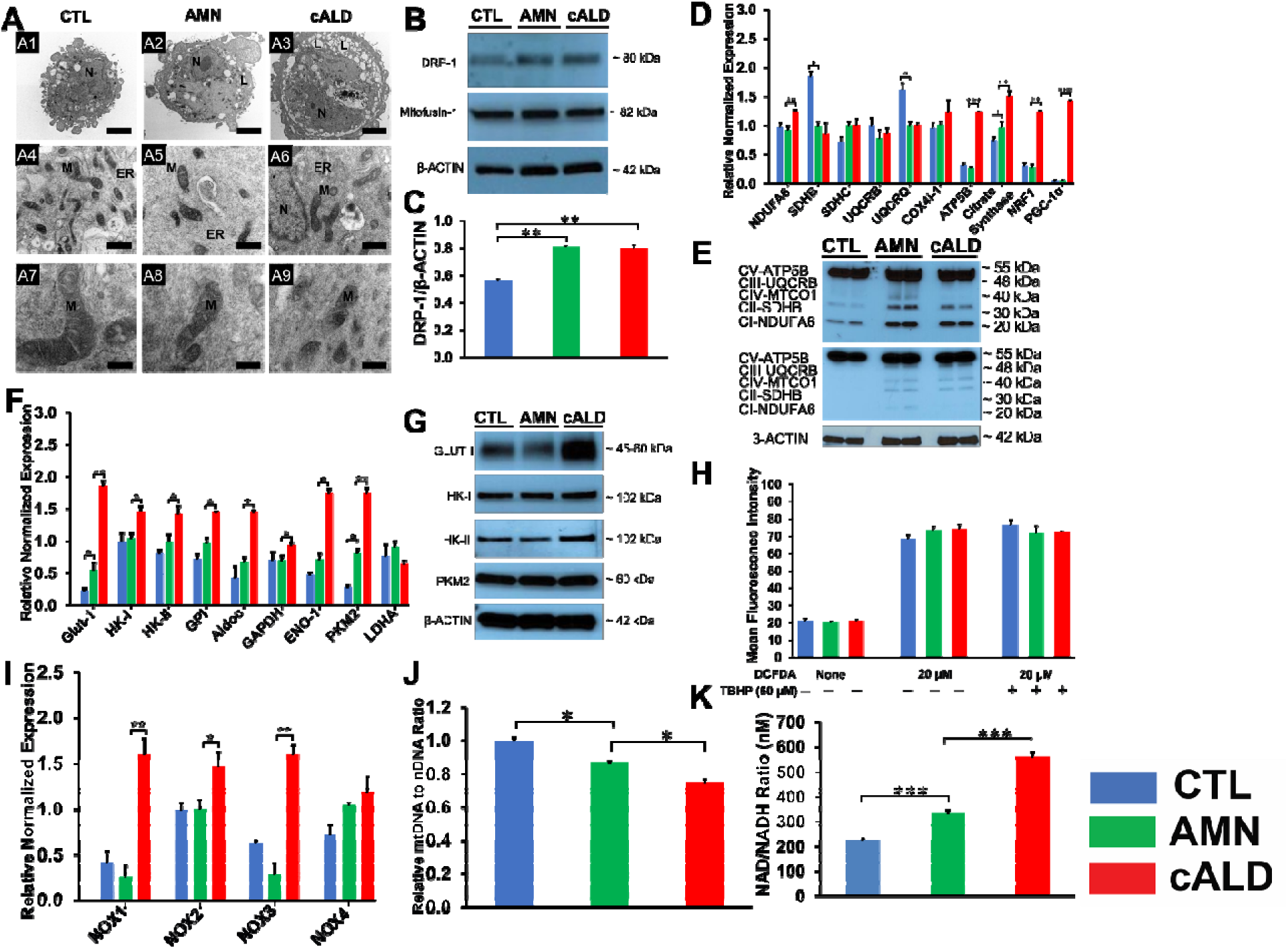
X-ALD patients show decreased mitochondrial content. Mitochondrial structural and functional assessment using TEM and RT-qPCR. (A) Representative TEM images of CTL, AMN, and cALD astrocytes. Upper panel (A1-A3 at scale bars 2 µm) represent whole CTL, AMN, and cALD astrocyte. Middle (A4-A6) and lower (A7-A9) represent mitochondrial structures in CTL, AMN, and cALD astrocytes (scale bars 1 µm and 400 nm, respectively). N: Nucleus; M: Mitochondria; ER: Endoplasmic reticulum. (B) Western blotting with DRP-1, mitofusin-1, and β-actin in CTL, AMN, and cALD patient astrocytes. (C) Quantification of DRP-1 (ratio against β-actin) in CTL, AMN, and cALD patient astrocytes (n = 3 technical replicates run in three independent blots). (D) RT-qPCR quantification of gene expression for mitochondrial genes (NDUFA6, SDHB, SDHC, UQCRB, UQCRQ, Cox4i-1, ATP5B, citrate synthase, and PGC1α) for CTL, AMN, and cALD astrocytes (n = 3 technical replicates per group). (E) Representative images for Western blotting with total OXPHOS at low exposure (bottom blot) and high exposure (top blot) and β-actin in CTL, AMN, and cALD patient astrocytes. (F) RT-qPCR quantification of gene expression for glycolytic genes (Glut-1, HK-I, HK-II, GPI, ALDOC, GapDH, EN-1, PKM2, and LDHA) for CTL, AMN, and cALD astrocytes (n = 3 technical replicates per group). (G) Representative images for Western blotting with Glut-I, HK-I, HK-II, PKM2, and and β-actin in CTL, AMN, and cALD patient astrocytes. (H) Quantification of reactive oxygen species (mean fluorescence intensity) between CTL, AMN, and cALD patient groups (n = 3 technical replicates per group). (I) RT-qPCR quantification of gene expression for NOX1, NOX2, NOX3, and NOX4 for astrocytes differentiated from CTL, AMN, and cALD iPSCs (n = 3 technical replicates per group). (J) Quantification of mtDNA to nDNA ratio in CTL, AMN, and cALD patient astrocytes (n = 3 technical replicates per group). (K) Quantification of NAD/NADH ratio in CTL, AMN, and cALD patient astrocytes (n = 3 technical replicates per group). **Data Information**: Data analysis for D, F, H, and I was performed using CFX Maestro software and the values represent mean means ± standard error. The gene expression of the individual sample was assessed with fold change using the comparative ΔΔ*Ct* method by normalizing with reference gene L-27 in CTL, AMN, and cALD astrocytes. The values in J are plotted as mtDNA/nDNA of AMN and cALD with reference ton CTL. Data in K represents corrected mean pixel density (pixel density – background) measured in 3 independent blots of DRP-1 on the samples (n = 3) and calculated as ratio of mean pixel density for DRP-1/β-actin. Unpaired two-tailed Student’s *t* test or Mann-Whitney test were used for analysis. Significance was denoted for P-values of less than 0.05. **P < 0.01; ***P < 0.001. **Abbreviations**: AMN: Adrenomyeloneuropathy; cALD: Cerebral adrenoleukodystrophy; CTL: Control; DRP-1: Dynamin related protein-1; mtDNA: Mitochondrial DNA; nDNA: Nuclear DNA; TEM: Transmission electron microscopy.

### Mitochondrial and Glycolytic Gene expressions are altered in AMN and cALD astrocytes

We performed RT-qPCR to assess if the altered OCR and ECAR in AMN and cALD astrocytes was also associated with alterations in expression of oxidative phosphorylation (OXPHOS) and glycolysis genes. Succinate dehydrogenase complex subunit B (SDHB) and ubiquinol-cytochrome C reductase complex ubiquinone-binding protein QP-C (UQCRQ) showed significantly lower expression in AMN and cALD astrocytes (p < 0.01 and p < 0.05, respectively; Figure 4D), suggesting defective transfer of electrons from NADH and succinate and possibly compromised inner membrane electrochemical gradient. cALD astrocytes showed significantly higher expression of NDUFA6 (NADH:Ubiquinone Oxidoreductase Subunit A6), ATP5B, citrate synthase, nuclear respiratory factor 1, and peroxisome proliferator-activated receptor gamma coactivator 1-α (PGC1α) than AMN astrocytes (p < 0.01; Figure 4D). AMN astrocytes showed higher expression of citrate synthase and PGC1α than CTL astrocytes (both p < 0.05; Figure 4D). Immunoblot analysis revealed less SDHB (Complex II-SDHB) and mitochondrially encoded cytochrome C oxidase I (Complex IV-MTCO1) in cALD astrocytes than in AMN astrocytes, whereas AMN astrocytes had more SDHB and MTCO1 than CTL astrocytes (Figure 4E).

In line with higher ECAR levels, cALD astrocytes also showed higher expression of glycolytic genes Glut-1/SLC2A, HK-I, HK-II, GPI, ALDOC, GAPDH, ENO-1, and PKM2 than AMN and CTL astrocytes, whereas LDHA was not significantly different among the three groups. AMN astrocytes did not show significant differences in expression of these genes relative to CTL astrocytes, except for PKM2 and GLUT1 which were significantly increased in AMN astrocytes ((Glut-1 and PKM2, p < 0.01; other genes, p < 0.01; Figure 4F and G).

### NADPH oxidase expression is increased in cALD astrocytes

Oxidative damage of proteins in observed in postmortem cALD brain and VLCFA accumulation is documented to induce oxidative stress. We measured the levels of ROS in CTL, AMN and cALD astrocytes. ROS generation did not significantly differ between the CTL, AMN and cALD groups under basal conditions (Figure 4H). However, NADPH oxidase (NOX) isoforms NOX1, NOX2 and NOX3 expression was significantly increased in cALD astrocytes (NOX1 and NOX3, p < 0.01; NOX2, p < 0.05; Figure 4I).

### AMN and cALD Astrocytes Show Reduced mtDNA Content

To determine if functional mitochondrial changes were associated with changes in mitochondrial content, we measured the mtDNA/nDNA ratios by RT-qPCR. mtDNA/nDNA ratio in AMN astrocytes was significantly lower (p<0.05) than CTL astrocyte (Figure 4J). cALD astrocytes had further significantly lower (p<0.05) mtDNA/nDNA ratio compared to AMN astrocytes suggesting lower mitochondrial content in X-ALD cells (p < 0.05; Figure 4J).

### Intracellular NAD/NADH Levels are Increased in AMN and cALD Astrocytes

NAD is a cofactor involved in electron transfer and NAD/NADH ratio reflects the intracellular redox state. cALD astrocytes showed a significantly higher NAD/NADH ratio (p < 0.001) than AMN astrocytes, which in turn had a significantly higher NAD/NADH ratio (p < 0.001) than CTL astrocytes (Figure 4K).

### Proinflammatory, Anti-inflammatory, Chemokine and neurotrophic Gene Expressions

We and others previously documented increased proinflammatory gene expression in postmortem brain tissue from patients with cALD (Paintlia AS, 2003; Singh *et al*., 2016). VLCFA accumulation is also associated with the inflammatory response in *ABCD1*-silenced mouse astrocytes (Singh *et al*., 2009). Therefore, we analyzed proinflammatory gene expression in unstimulated AMN and cALD iPSC-derived astrocytes. Astrocytes from cALD patients showed significantly higher interferon-γ (p < 0.01), nitric oxide synthase 2, TNF-α (both p < 0.05), IL-17A, and IL-22 (both p < 0.01) expression than AMN astrocytes (Figure 5A). TNF-α expression in AMN was also significantly higher than CTL astrocytes (p < 0.05; Figure 5A). IL-12α (p < 0.01) and IL-12β (p < 0.05) expression was lower in cALD astrocytes compared with AMN and CTL astrocytes (Figure 5A). IL-12α expression in AMN was comparable to CTL but IL-12β was significantly decreased compared to CTL astrocytes (p < 0.01; Figure 5A). AMN astrocytes had higher IL-6 expression than cALD and CTL astrocytes (p < 0.01; Figure 5A). IL-1β (p < 0.05) and IL-RA (p < 0.01) showed lower expression in both AMN and cALD astrocytes compared with CTL (Figure 5A), although IL-1R was significantly higher in AMN and cALD astrocytes than in CTL astrocytes (p < 0.05). IL-1β and IL-23α expression was comparable between AMN and cALD astrocytes (Figure 5A).

**Figure 5:**
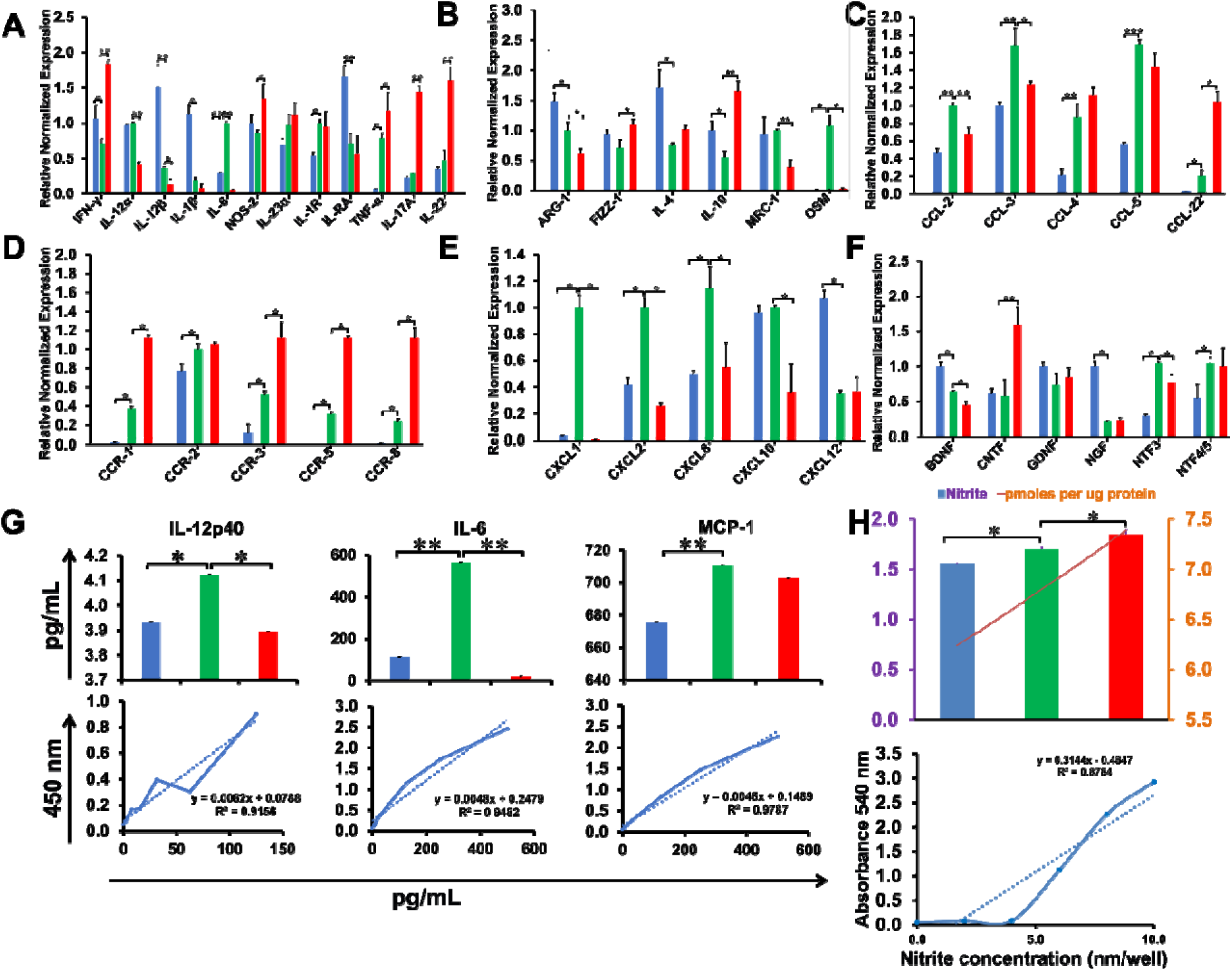
cALD astrocytes reveal an increased proinflammatory cytokine expression and a higher nitrite concentration. (A & B) RT-qPCR quantification of gene expression for proinflammatory (IFN-γ, IL-12α, IL-12β, IL-1β, IL-6, Nos2, IL-23α, IL-1R, IL-RA, TNF-α, IL-17A, and IL-22) (A) and anti-inflammatory (Arg-1, Fizz-1, IL-4, IL-10, MRC-1, and OSM) (B) in CTL, AMN, and cALD astrocytes (n = 3 technical replicates per line for astrocytes). The gene expression of the individual sample was assessed with fold change using the comparative ΔΔ*Ct* method by normalizing with reference gene L-27 in CTL, AMN, and cALD astrocytes. (C-E) RT-qPCR quantification of gene expression for C-C motif chemokine ligands (CCL-2, CCL-3, CCL-4, CCL-5, and CCL-22) (C), C-C motif chemokine receptors (CCR-1, CCR-2, CCR-3, CCR-5, and CCR-8) (D), and C-X-C motif chemokine ligands (CXCL1, CXCL2, CXCL8, CXCL10, and CXCL12) (E) in CTL, AMN, and cALD astrocytes (n = 3 technical replicates per line for astrocytes). (F) RT-qPCR quantification of gene expression for neurotrophic factors (BDNF, CNTF, GDNF, NGF, NTF3, and NTF4/5) for CTL, AMN, and cALD astrocytes (n = 3 technical replicates per line for astrocytes). The gene expression of the individual sample was assessed with fold change using the comparative ΔΔ*Ct* method by normalizing with reference gene L-27 in CTL, AMN, and cALD astrocytes. (G) Quantification of IL-12p40, IL-6, and MCP-1 in the culture supernatant of CTL, AMN, and cALD astrocytes. Results represent means ± standard error (pg/mL) from triplicates of each sample type (n = 3). (H) Quantification of nitrites in CTL, AMN, and cALD astrocyte lysates. Panel H represents the nitrites (OD and pmoles per µg protein) calculated using standard curve in the panel C. Color scheme shows colors assigned to sample types in qRT-PCR plots. **Data information**: Results represent means ± standard error from triplicates of each sample type. Two-tailed Student’s *t* test or Mann-Whitney tests were performed to analyze the data. Significance was denoted for P values as *P < 0.05, **P < 0.01; ***P < 0.001. **Abbreviations**: AMN: Adrenomyeloneuropathy; cALD: Cerebral adrenoleukodystrophy; CTL: Control; OD: Optical density (absorbance).

On the other hand, anti-inflammatory cytokines arginase-1 (ARG-1) and mannose receptor C-type-1 expression was significantly lower in cALD astrocytes than in AMN astrocytes (both p < 0.05; Figure 5B). ARG-1 in AMN astrocytes was in turn significantly lower than CTL astrocytes (p < 0.05). CTL astrocytes had significantly higher IL-4 levels compared to both AMN and cALD astrocytes (p < 0.05). IL-4 levels between AMN and cALD were not significantly different. Fizz-1 expression was higher only in cALD astrocytes relative to CTL astrocytes (p < 0.05). IL-10 expression was significantly lower (p < 0.05) in AMN astrocytes than CTL astrocytes and was significantly higher (p < 0.01) in cALD astrocytes than in AMN astrocytes. Oncostatin M (OSM) was significantly higher in AMN compared to CTL and cALD astrocytes (p < 0.05; Figure 5B).

CCL-2 and other chemokines play key roles in the pathogenesis of neurological disorders, including X-ALD (Paintlia AS, 2003; Weinhofer et al., 2018). Thus, we analyzed chemokine expression in unstimulated CTL, AMN and cALD iPSC-derived astrocytes. CCL-2, CCL-3, CCL-4 (all p < 0.01), CCL5 (p < 0.001) and CCL-22 (p < 0.05) expression were significantly higher in AMN than in CTL astrocytes (Figure 5C). In cALD astrocytes, CCL-2 (p < 0.01) and CCL-3 (p < 0.05) were significantly lower and CCL-22 (p < 0.05) was significantly higher than in AMN astrocytes (Figure 5C). AMN astrocytes had significantly higher expression of chemokine receptors CCR-1, CCR-2, CCR-3, CCR-5, and CCR-8 than CTL astrocytes (all p < 0.05; Figure 5D). Expression of these chemokine receptors was further significantly higher in cALD than in AMN astrocytes (all p < 0.05; Figure 5D). CXCL-1, CXCL-2, and CXCL-8 expression were significantly higher (p < 0.05) in AMN astrocytes than in CTL astrocytes and showed lower expression in cALD astrocytes than in AMN astrocytes (p < 0.05; Figure 5E). AMN and CTL astrocytes had comparable expression of CXCL10, while cALD astrocytes had significantly lower expression of CXCL10 than AMN and CTL astrocytes (p < 0.05). Both AMN and cALD astrocytes had significantly lower expression of CXCL12 than CTL astrocytes (p < 0.05; Figure 5E).

Astrocytes are a major source of neurotrophic support in the central nervous system (Ricci et al., 2009). We next investigated if neurotrophic gene expressions were altered between CTL, AMN, and cALD astrocytes. AMN astrocytes showed significantly lower expression of BDNF compared with CTL astrocytes (p < 0.05), and BDNF expression was further significantly decreased in cALD astrocytes (p < 0.05; Figure 5F). CNTF expression was significantly higher in cALD (p < 0.01) than in AMN and CTL astrocytes, and no significant change in GDNF levels were seen between the three astrocyte groups. Both AMN and cALD astrocytes had significantly lower expression of NGF (p < 0.05) and significantly higher expression of NTF3 and NTF4/5 (both p < 0.05) compared with CTL astrocytes (Figure 5F).

### Enzyme-Linked Immunosorbent and Nitrite Assay

AMN astrocytes contained significantly higher levels of IL-12p40 than CTL and cALD astrocytes (p < 0.05; Figure 5G). Levels of IL-12p40 were not significantly different between CTL and cALD astrocytes. Both IL-6 and MCP-1 were significantly higher in AMN astrocytes than in CTL and cALD astrocytes (both p < 0.01; Figure 5G). IL-6 in cALD astrocytes was significantly lower than in CTL astrocytes (p < 0.05), whereas TNF-α and IL-1β were undetectable in culture supernatants of all astrocytes (data not shown). We assessed intracellular nitric oxide and observed that basal levels of nitric oxide were significantly higher (p < 0.05) in AMN astrocytes than in CTL astrocytes and was further significantly increased in cALD astrocytes (p < 0.05; Figure 5H).

### cALD Astrocytes have Increased STAT3 (Serine^727^) Phosphorylation and Elevated Toll-Like Receptor Expression

To determine the cell signaling underlying the differential cytokine and chemokine response in AMN and cALD astrocytes, we used a human phospho-kinase array proteome profiler to identify inflammatory signaling. STAT3 phosphorylation at Serine^727^ was significantly higher in cALD astrocytes (Figure 6A), which was confirmed by densitometry (p < 0.001; Figure 6B) and immunoblot analysis (Figure 6C). We identified hyperphosphorylation of Akt, mitogen-activated protein kinase (MAPK) p38, Ras-dependent extracellular signal-regulated kinase (ERK) 1/2, and stress-activated protein kinases/Jun amino-terminal kinases, as well as reduced phosphorylation of adenosine monophosphate-activated protein kinase α1 (AMPKα1) and β-catenin in cALD astrocytes (Figure 6D-E).

**Figure 6:**
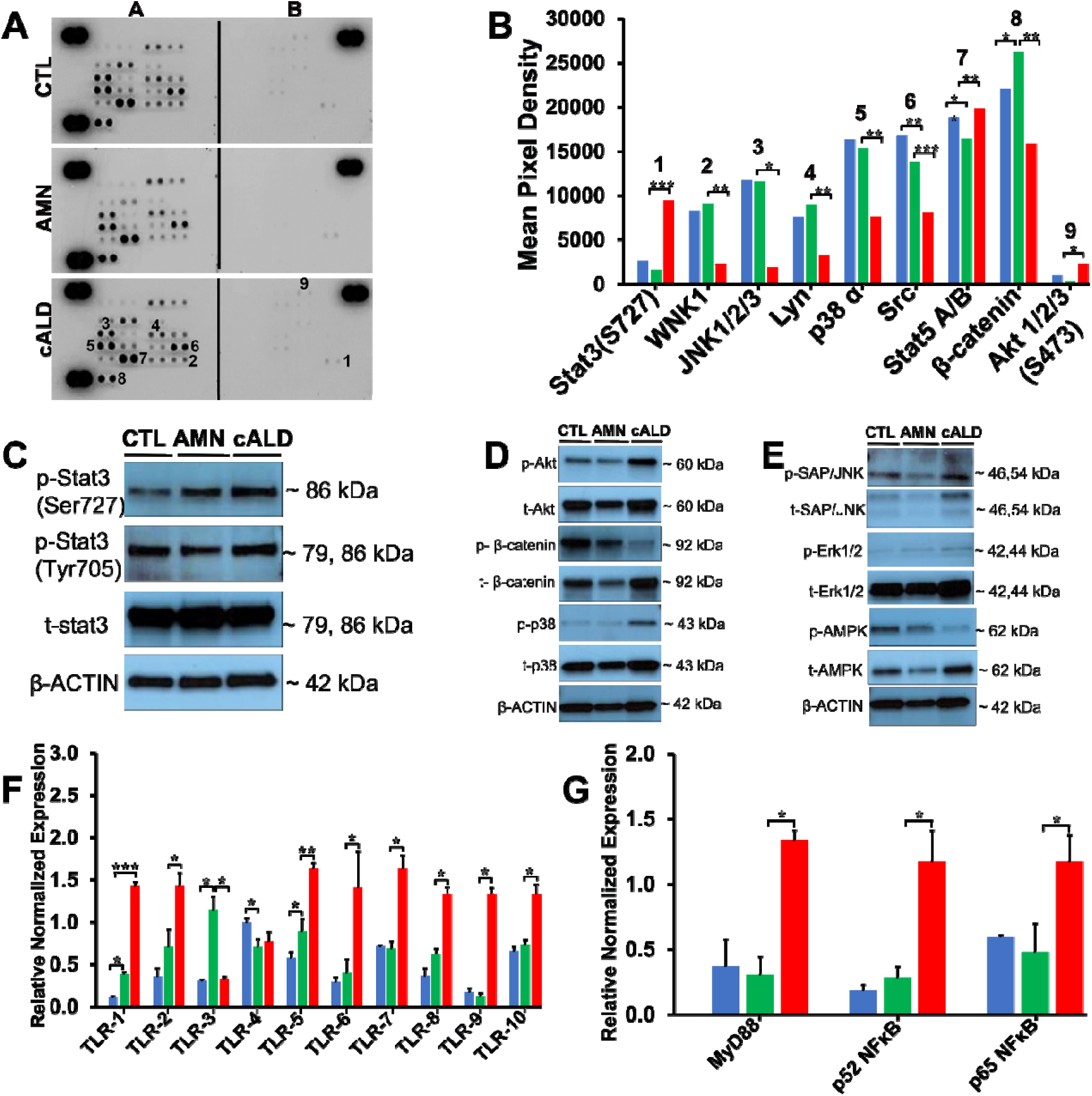
cALD astrocytes show higher Stat3 and TLR activity. (A & B) Phosphokinase antibody array blots using CTL, AMN, and cALD patient astrocyte lysates (250 µg for each of antibody array membranes) (A) and Kinase phosphorylation analysis using mean pixel density showing basal level changes in the expression levels of Stat3, JNK1/2/3, p38 α, SRC, Stat5 A/B, β-catenin, and Akt1/2/3 activity in CTL, AMN, and cALD astrocytes (B). (C) Representative images for Western blotting with p-Stat3 (ser727), p-Stat3 (tyr705), total Stat3, and β-actin in CTL, AMN, and cALD patient astrocytes. (D & E) Representative images for Western blotting with phosphorylated and total Akt, β-catenin, and p38 (D) and SAP/JNK, Erk1/2, and AMPK (E) in CTL, AMN, and cALD patient astrocytes. β-actin was used as reference CTL. (F & G) RT-qPCR quantification of gene expression for TLR (TLR1-10) (F) and MyD88, p52 NFκB, and p65 NFκB (G) for CTL, AMN, and cALD astrocytes (n = 3 technical replicates per line for astrocytes). The gene expression of the individual sample was assessed with fold change using the comparative ΔΔ*Ct* method by normalizing with reference gene L-27 in CTL, AMN, and cALD astrocytes. Color scheme shows colors assigned to sample types in qRT-PCR plots. **Data information**: Results represent means ± standard error from triplicates of each sample type. Two-tailed Student’s *t* test or Mann-Whitney tests were performed to analyze the data. Significance was denoted for P-values of less than 0.05. **P < 0.01; ***P < 0.001. **Abbreviations**: AMN: Adrenomyeloneuropathy; cALD: Cerebral adrenoleukodystrophy; CTL: Control; Stat3: Signal transducer and activator of transcription 3; TLR: Toll-like receptor.

We interrogated astrocytes for toll-like receptor (TLR) expression. AMN astrocytes showed higher expression of TLR-1, TLR-3, and TLR-5 than CTL astrocytes (all p < 0.05; Figure 6F). TLR-4 levels were significantly lower in AMN astrocytes than in CTL astrocytes (p < 0.05; Figure 6G). Interestingly, cALD astrocytes had significantly higher expression of TLR-1 (p < 0.001), TLR-2 (p < 0.05), TLR-5 (p < 0.01), TLR-6 (p < 0.05), TLR-7 (p < 0.05), TLR-8 (p < 0.05), TLR-9 (p < 0.05), and TLR-10 (p < 0.01) than AMN astrocytes. We also observed significantly elevated expression of the major TLR adaptor protein myeloid differentiation factor 88 (MyD88) in cALD astrocytes (p < 0.05; Figure 6G). Furthermore, we observed significantly higher expression of NF-κB subunits p52 (NF-κB2) and p65 (RelA) in cALD astrocytes than in AMN astrocytes (p < 0.05; Figure 6G).

### MicroRNA Sequencing and Analysis

MicroRNA (miRNA) post-transcriptionally regulate gene expression. To comprehensively screen differentially expressed miRNA in unstimulated CTL, AMN and cALD iPSC-derived astrocytes, we performed profiling for miRNA using miRNA-Seq. We identified 12 miRNAs (Wald test, False Discovery Rate < 0.05) that were differentially expressed between cALD, AMN, and CTL astrocytes (Figure 7A-B). There were eight miRNAs that were more abundant and three that showed lower expression in AMN astrocytes relative to CTL astrocytes. The cALD astrocytes showed higher expression of three miRNAs relative to CTL astrocytes, although the expression levels were lower than in the AMN cells. Notably, hsa-miR-9-3p and hsa-miR-9-5p demonstrated a phenotype severity-specific directional change with highest expression in cALD (CTL<AMN<cALD; Figure 7B).

**Figure 7:**
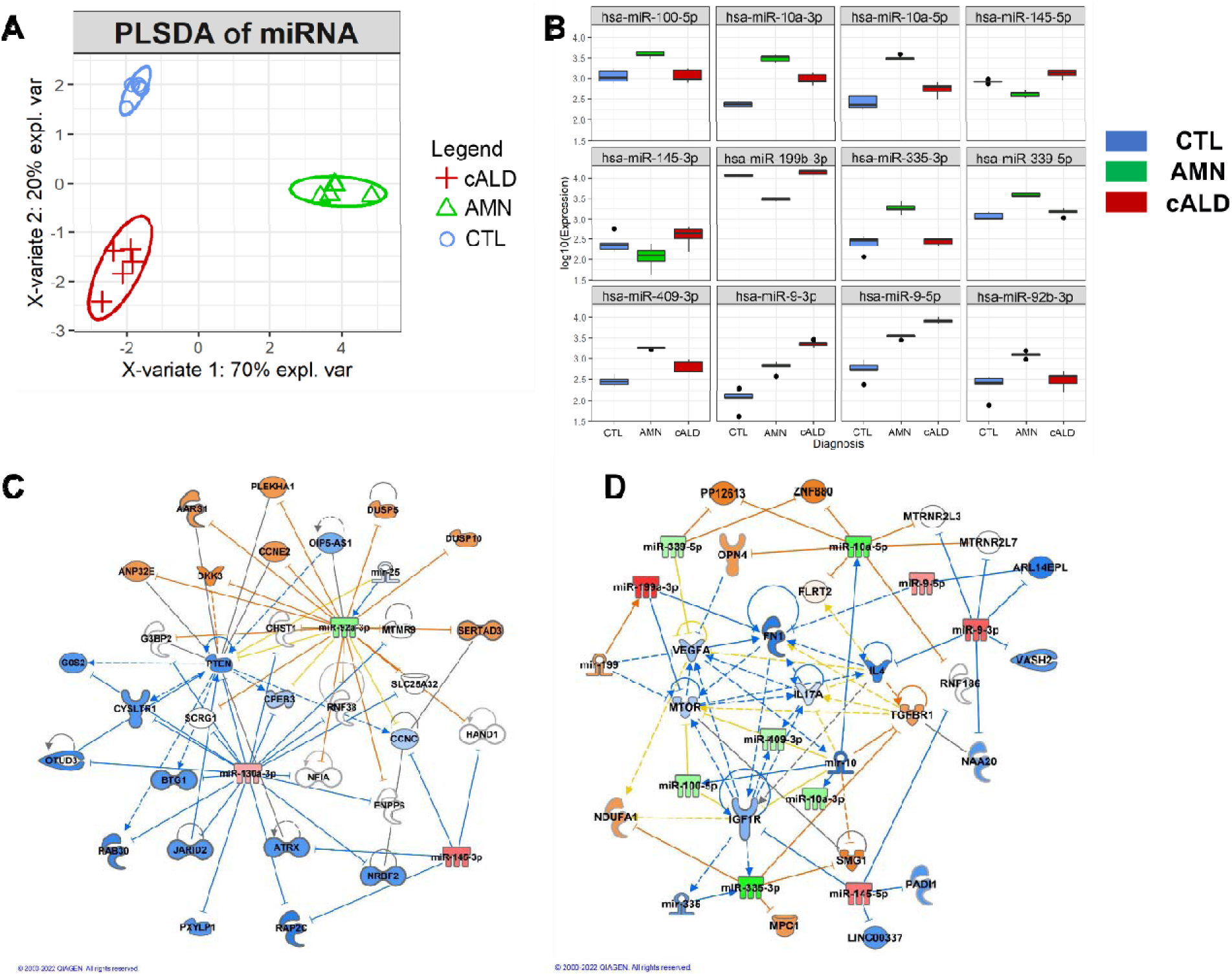
CTL, AMN, and cALD patient astrocytes retain biochemical characteristics of X-ALD patients. (A) Partial least squares discriminant analysis (PLSDA) score plots of significant miRNAs determined by miRNA sequence analysis of CTL, AMN, and cALD patient astrocytes (n = 5 technical replicates of each sample type). (B) Box plots of 12 significant miRNAs determined by miRNA sequencing of CTL, AMN, and cALD iPSC-derived astrocytes. (n = 5 technical replicates of each sample type). (C & D) Ingenuity pathway gene network generated by comparing differentially expressed miRNAs and regulatory pathways in CTL vs AMN, and AMN vs cALD astrocytes. **Data information:** Differential miRNAs in A were identified using Wald Test. False discovery was controlled with Benjamini-Hochberg corrected *P*-values, with adjusted-p < 0.05 and mean fold change >2 as the threshold for statistical significance. Boxplots in B represent the miRNA abundance (log10) within the three groups, for commonly altered miRNAs. C and D are networks of interrelated miRNA and associated molecular components generated using QIAGEN IPA. **Abbreviations**: AMN: Adrenomyeloneuropathy; cALD: Cerebral adrenoleukodystrophy; CTL: Control; miRNA: microRNA; PLSDA: Partial Least Square-Discriminant Analysis

To examine biological significance of differentially expressed miRNA between CTL and AMN (Figure 7C) and between AMN and cALD astrocytes (Figure 7D), pathway enrichment was conducted using the ingenuity pathway analysis (IPA). Between CTL and AMN astrocytes, the central pathways associated with altered miRNA were PTEN, ATRX, and HAND1 (Figure 7C) which are associated with neuronal maturation and gliomas (Alghamri et al., 2021; Kang et al., 2020). On the contrary, central to miRNA altered between AMN and cALD astrocytes were pathways associated with immune function regulated through IGF1R, IL-17A, and mTOR (Figure 7D).

### CRISPR-Cas9 knock-in of *ABCD1* in AMN and cALD Astrocytes

We next investigated if knock-in of a functional *ABCD1* in AMN and cALD astrocytes may reverse of the aberrant molecular signature identified above. *ABCD1* gene was delivered using a Lentiviral vector in AMN and cALD astrocytes successfully expressed *ABCD1* protein in AMN (AMN-OV) and cALD (cALD-OV) astrocytes. *ABCD1* expression did not seem to have off-target effects since the level of ABCD2 and ABCD3 protein remained the same (Figure 8A). In OXPHOS analysis by immunoblot, we observed that mitochondrial subunit I NDUFA6 and subunit II SDHB were upregulated in AMN-OV and cALD-OV compared to the AMN and cALD astrocytes respectively, whereas the other subunits did not show a change after *ABCD1* knock-in (Figure 8B). STAT3 phosphorylation at Serine^727^ and Tyrosine^705^ was increased in AMN-OV and cALD-OV indicating that *ABCD1* induction did not reverse the hyperphosphorylation (Figure 8C). We assessed basal gene expression profile and effect of *ABCD1* knock-in on their expression in AMN and cALD astrocytes. Induction of *ABCD1* gene expression significantly increased SDHB (p < 0.05), SDHC (p < 0.05), UQCRQ (p < 0.05), ATP5B (p < 0.01), NRF1 (p < 0.01), and PGC-1α (p < 0.01) in AMN-OV. However, *ABCD1* activation reduced expression of NRF1 and PGC-1α in cALD-OV astrocytes (both p < 0.05; Figure 8D). We did not observe a change in NDUFA6, UQCRB, Cox4i1, and Citrate Synthase in AMN-OV and cALD-OV astrocytes (Figure S1). Glycolytic genes Aldoc and ENO-1 showed decreased expression in AMN-OV astrocytes (both p < 0.05; Figure 8E). Glut-1, HK-I, HK-II, GPI, Aldoc, ENO-1, PKM2, and LDHA which showed higher expression in cALD astrocytes were reduced in expression upon *ABCD1* knock-in in cALD-OV astrocytes (ENO-1, p < 0.01; all others p < 0.05; Figure 8E). However, Glut-1 and GAPDH were upregulated in AMN-OV astrocytes (both p < 0.05; Figure 8D and S1). *ABCD1* induction significantly mitigated the expression of NOX1 and NOX2 in cALD-OV (both p < 0.01; Figure 8F). On the other hand, NOX3 and NOX4 increased upon *ABCD1* induction in cALD-OV astrocytes (both p < 0.05; Figure S1). Interestingly, *ABCD1* knock-in significantly lowered the expression of proinflammatory genes, IFN-y, IL-12α, NOS-2, and IL-23α in both AMN-OV and cALD-OV astrocytes whereas IL-1β expression was lowered in AMN-OV only. (all p < 0.05; Figure 8F). *ABCD1* knock-in resulted in increased expression of anti-inflammatory genes, Fizz-1 and OSM (both p < 0.05) in both AMN-OV and cALD-OV astrocytes; however, IL-4 (p < 0.05), IL-10, (p < 0.01) and MRC-1 (p < 0.05) were upregulated only in AMN-OV astrocytes. IL-4 decreased (p < 0.01) in cALD-OV and IL-10 and MRC-1 remained unchanged in cALD-OV astrocytes (Figure 8H). There was no change in IL-12 β, IL-6, IL-1R, IL-RA, and TNF-α expression in AMN and cALD after *ABCD1* knock-in (Figure S1). IL-17A (p < 0.01) and IL-22 (p < 0.05) showed significant increase in AMN-OV astrocytes, however, IL-17A was significantly reduced in cALD-OV (p < 0.05) astrocytes whereas IL-22 did not change in cALD-OV astrocytes after *ABCD1*knock-in (Figure S1). We observed increase in CCL-3 (p < 0.05) and CCR-3 (p < 0.01) chemokines in AMN-OV astrocytes, whereas CCR-2 (p < 0.01) and CXCL12(p < 0.05) increased in both AMN-OV and cALD-OV astrocytes upon *ABCD1* induction. CCR-5 (p < 0.01) and CCR-8 (p < 0.05) increased in AMN-OV whereas decreased in cALD-OV astrocytes (p < 0.05) (Figure 8I). We did not observe a change in expression of other CCLs (CCL-4, -5, and -22), CCR-1, and CXCLs (CXCL-1, -2, -8, and -10) after *ABCD1* knock-in (Figure S1). *ABCD1* knock-in resulted in significantly increased expression of BDNF (p < 0.01), CNTF (p < 0.05), and NGF (p < 0.05) in AMN-OV astrocytes whereas NGF was upregulated in cALD-OV astrocytes (p < 0.01). On the other hand, CNTF showed significant downregulation (p < 0.05) and BDNF was unchanged in cALD-OV astrocytes (Figure 8J). We did not any significant change in expressions of GDNF, NTF3, and NTF4/5 in AMN-OV and cALD-OV astrocytes after *ABCD1* knock-in (Figure S1). TLR 1-2, and TLR 5-10 expression were significantly reduced in cALD-OV astrocytes (TLR-5 and TLR-7 p < 0.01; TLR-1,-2, -6, 8-10, p < 0.05); however, but intriguingly increased in AMN-OV astrocytes. TLR-3 and TLR-4 remained unchanged (Figure 8K and S1). Hsa-miR-9-3p and hsa-miR-9-5p were increased with the severity of disease phenotype with highest expression in cALD astrocytes and lowest in CTL(CTL<AMN, p < 0.01; AMN < cALD p < 0.05). *ABCD1* knock-in of *ABCD1* significantly decreased the expression of miR-9-3p (p < 0.001) and miR-9-5p (p < 0.01) in AMN-OV astrocytes but not in cALD-OV astrocytes (Figure 8L).

**Figure 8:**
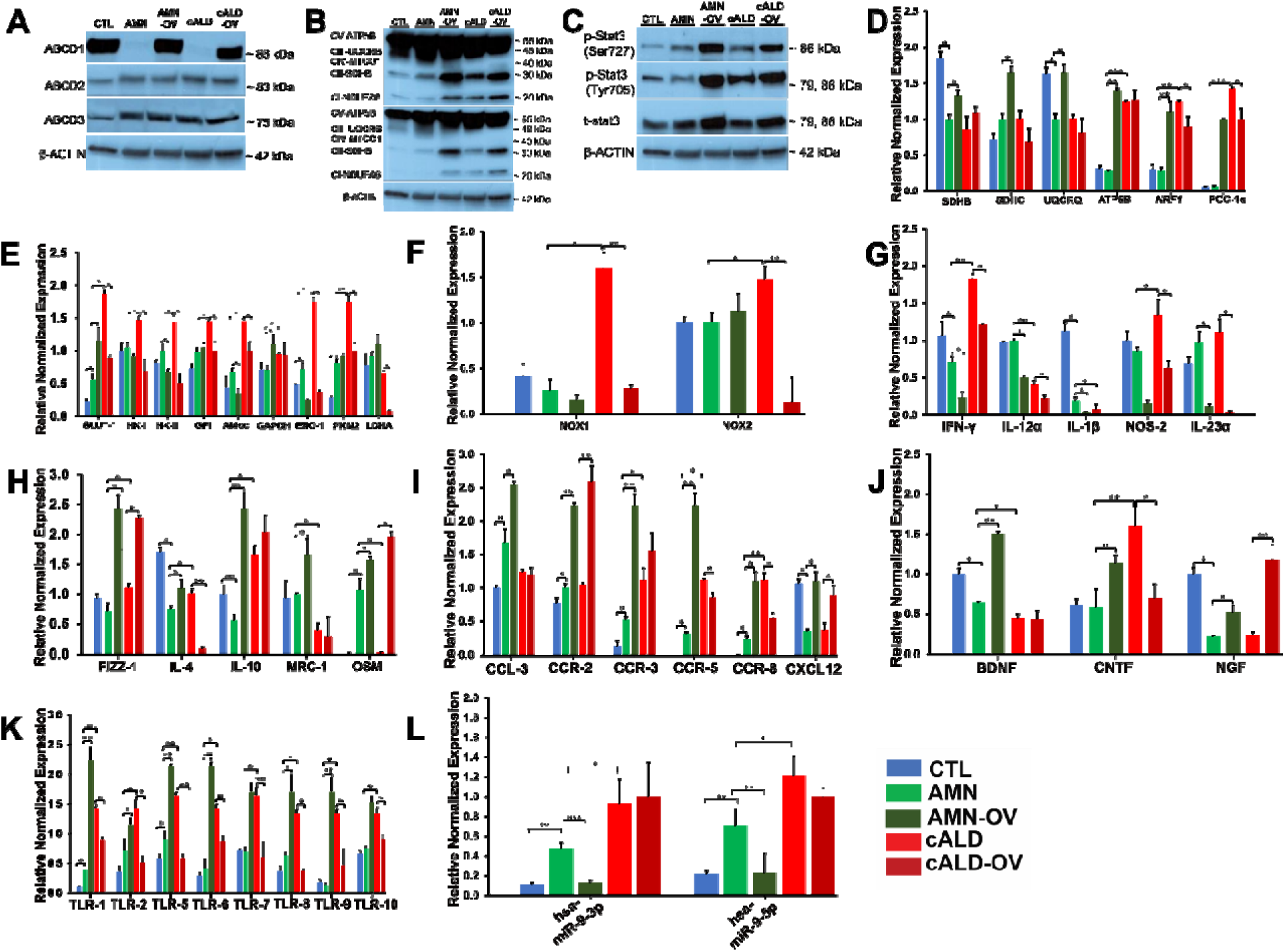
*ABCD1* Gene Induction Lowers Inflammation in AMN and cALD Astrocytes. (A) Representative images for Western blotting with *ABCD1*, ABCD2, and ABCD3, and β-actin in CTL, AMN, AMN-OV, cALD, and cALD-OV patient astrocytes. (B) Representative images for Western blotting with total OXPHOS at low exposure (bottom blot) and high exposure (top blot) and β-actin in CTL, AMN, AMN-OV, cALD, and cALD-OV astrocytes. (C) Representative images for Western blotting with p-Stat3 (ser727), p-Stat3 (tyr705), total Stat3, and β-actin in CTL, AMN, AMN-OV, cALD, and cALD-OV astrocytes. (D & E) RT-qPCR quantification of gene expression for mitochondrial genes (D) and glycolytic genes (E) for CTL, AMN, AMN-OV, cALD, and cALD-OV astrocytes (n = 3 technical replicates per line for astrocytes). (F) RT-qPCR quantification of gene expression for NADPH Oxidases for CTL, AMN, AMN-OV, cALD, and cALD-OV astrocytes (n = 3 technical replicates per line for astrocytes). (G & H) RT-qPCR quantification of gene expression for proinflammatory (IFN-γ, IL-12α, Nos2, and IL-23α) (G) and anti-inflammatory (Fizz-1, IL-4, IL-10, and MRC-1) (H) in CTL, AMN, AMN-OV, cALD, and cALD-OV astrocytes (n = 3 technical replicates per line for astrocytes). The gene expression of the individual sample was assessed with fold change using the comparative ΔΔ*Ct* method by normalizing with reference gene L-27 in CTL, AMN, and cALD astrocytes. (I) RT-qPCR quantification of gene expression for C-C motif chemokine ligands (CCL-3), C-C motif chemokine receptors (CCR-2, CCR-3, CCR-5, and CCR-8), and C-X-C motif chemokine ligands (CXCL12) in CTL, AMN, AMN-OV, cALD, and cALD-OV astrocytes (n = 3 technical replicates per line for astrocytes). (J) RT-qPCR quantification of gene expression for neurotrophic factors (BDNF, CNTF, and NGF) for CTL, AMN, AMN-OV, cALD, and cALD-OV astrocytes (n = 3 technical replicates per line for astrocytes). The gene expression of the individual sample was assessed with fold change using the comparative ΔΔ*Ct* method by normalizing with reference gene L-27 in CTL, AMN, and cALD astrocytes. (K) RT-qPCR quantification of gene expression for TLR (TLR-1, -2, & 5-10) in CTL, AMN, AMN-OV, cALD, and cALD-OV astrocytes (n = 3 technical replicates per line for astrocytes). The gene expression of the individual sample was assessed with fold change using the comparative ΔΔ*Ct* method by normalizing with reference gene L-27 in CTL, AMN, and cALD astrocytes. Color scheme shows colors assigned to sample types in qRT-PCR plots. (L) RT-qPCR quantification of gene expression for hsa-miR-9-3p and hsa-miR-9-5p in CTL, AMN, AMN-OV, cALD, and cALD-OV astrocytes (n = 3 technical replicates per line for astrocytes). The gene expression of the individual sample was assessed with fold change using the comparative ΔΔ*Ct* method by normalizing with reference miRNA RNU6 in CTL, AMN, and cALD astrocytes. **Data information:** Results represent means ± standard error from triplicates of each sample type. Two-tailed Student’s *t* test or Mann-Whitney tests were performed to analyze the data. Significance was denoted for P values as *P < 0.05, **P < 0.01; ***P < 0.001. Abbreviations: AMN: Adrenomyeloneuropathy; cALD: Cerebral adrenoleukodystrophy; CTL: Control; OD: Optical density (absorbance).

## DISCUSSION

To date, the mechanistic underpinnings of AMN and cALD phenotypes in X-ALD remain poorly understood. While others have used X-ALD human fibroblast-derived iPSCs to generate brain glial cells (Baarine et al., 2015; Jang et al., 2011), we report novel and comprehensive differences in the phenotypic profiles of cALD and AMN astrocytes. Consistent with previous reports, *ABCD1* variants in patient fibroblasts did not impede reprogramming and differentiation into astrocytes (Jang *et al*., 2011; Wang et al., 2012). iPSC-derived astrocytes from AMN and cALD phenotype not only showed characteristic X-ALD features, including *ABCD1* loss and higher VLCFA levels, but also displayed differential molecular and metabolic behavior. Heightened STAT3, TLR expression, cytokine response and novel increased miR-9 expression observed in cALD astrocytes may play a mechanistic role in differential neuroinflammatory response between AMN and cALD phenotypes.

Oxidative stress is considered an early even in X-ALD pathology. Accumulation of VLCFA causes oxidative stress, affects glycolysis and OXPHOS (Fourcade et al., 2008), and induces mitochondrial dysfunction (Singh and Giri, 2014; Singh *et al*., 2016; Singh *et al*., 2015). cALD astrocytes demonstrated increased glycolysis, glycolytic capacity, and elevated glycolytic gene expression, and hyperglycolysis in cALD astrocytes suggests a higher energy demand, likely making them more prone to inflammation than AMN astrocytes. While a hyperglycolytic state is implicated in neuroinflammation (Pamies D, 2021), activated STAT3 may lead to altered mitochondrial energetics (Wegrzyn et al., 2009). STAT3, a potent inducer of astrogliosis and astrocyte differentiation (Hong and Song, 2014) can be activated by either the Janus family of kinases or intrinsic tyrosine kinase activity from growth factors and cytokines (Shao et al., 2003) phosphorylating both serine^727^ and tyrosine^705^(Wen et al., 1995). cALD astrocytes increased phosphorylation of STAT3-p-serine^727^ and STAT3-p-tyrosine^705^ indicating that cALD astrocytes may be highly reactive. Furthermore, elevated neurotrophic factors such as ciliary neurotrophic factor in cALD may also signal through STAT3 and trigger reactive astrocytosis (Herrmann JE, 2008). STAT3 and CCL2 among the eight cluster genes were identified to share expression patterns in cALD and Alzheimer’s disease; however, gene correlation network analysis pattern of AMN and Alzheimer’s disease is not available (Shim YJ, 2022). Higher CCL2 observed in AMN astrocytes maybe protective by virtue of its role in microglial recruitment and regulating the immune system (Ueno et al., 2000).

Metabolic profiling indicated a higher non-mitochondrial OCR, which is associated with enzymes associated with inflammation including NADPH oxidases (Chacko et al., 2014) and a higher maximal OCR which may be attributed to higher electron transport capacity in cALD astrocytes (Chacko *et al*., 2014). Similarly, we observed an increased non-glycolytic acidification, higher glycolytic reserve, and glycolytic capacity in cALD astrocytes which may be due to increased uncoupling and mitochondrial dysfunction. Increased DRP-1 expression in cALD than in AMN astrocytes, suggesting higher mitochondrial fission, and a significant decrease in the mtDNA/nDNA ratio in AMN and cALD astrocytes suggests an overall decreased mitochondrial content. Accumulation of dysfunctional mitochondria can lead to ROS production. Although, we did not detect increased ROS in cALD astrocytes, expression of superoxide catalyzing NADPH oxidases and NDUFA6 were increased. We also observed reduced SDHB (complex II) protein in both AMN and cALD astrocytes, implying impaired electron transport. These data suggest that activated STAT3 may play a role in inflammation in cALD astrocytes, altering mitochondrial OCR.

mtDNA released by damaged mitochondria due to activated STAT3 (Wegrzyn *et al*., 2009) may induce endogenous TLR signaling, especially through TLR9. *In vitro*, TLR9 antagonism modulates astrocyte chemokine release and macrophage chemotaxis in spinal cord injury (Li et al., 2020). In line with this, we observed higher expression of TLR1-2, TLR5-10, MyD88, and NF-κB expression in cALD astrocytes. This is also consistent with upregulated TLR-MyD88-NF-κB pathway in the spinal cord of *ABCD1*-KO mice and increased NF-κB levels in *ABCD1*-KO mice astrocytes (Schluter et al., 2012; Singh *et al*., 2009). Furthermore, upregulated Akt and p-Akt in cALD astrocytes implies that TLR in association with p-Akt signaling may further trigger reactive astrocytes in patients with cALD (Cheng et al., 2020). Despite lower relative mitochondrial content and higher DRP-1 expression in AMN relative to CTL astrocytes, AMN astrocytes had decreased STAT3 (serine^727^) phosphorylation, lower NOX and TLR expression, which may underlie the milder non-inflammatory phenotype in AMN.

Increased NAD/NADH ratio is associated with disrupted energy homeostasis, and we observed significantly higher intracellular NAD/NADH ratios in both AMN and cALD astrocytes. A previous study reported a reduced NAD/NADH ratio in *ABCD1*-silenced human U87 cells (Baarine M, 2015). This discordance may be due to the nature of the source of the U87 cells, which are derived from a human glioma patient, compared to AMN and cALD patient-iPSC-derived astrocytes employed in the present study. Moreover, higher NAD/NADH ratios observed in AMN and cALD astrocytes could be due to higher glycolysis, where NADH is oxidized to NAD^+^ and the excess NADH is shuttled to mitochondria where NAD^+^ is reduced to NADH (Yang and Sauve, 2016). Mitochondrial dysfunction as seen in cALD patients, may result in impaired reduction of NAD^+^ to NADH in mitochondria.

We previously documented loss of AMPKα1 in cALD fibroblasts, glial cells, and cALD postmortem brain white matter (Singh and Giri, 2014; Singh *et al*., 2016; Singh *et al*., 2015). AMPKα1 is the master regulator of energy homeostasis, and its perturbation is associated with neuroinflammatory diseases (Nath et al., 2009). Induction of glycolysis is promoted by growth factors via signal transduction mediated by the Akt pathway (Fox et al., 2005), whereas AMPKα1 is an inhibitor of the metabolic switch from OXPHOS to aerobic glycolysis. Consequently, a higher p-Akt and loss of AMPKα1 in cALD astrocytes can explain the hyper-glycolytic state of cALD astrocytes. Concomitant with the anti-inflammatory role of AMPKα1, we observed significant upregulation of cytokine and chemokine response in cALD astrocytes. The Th17-linked cytokine IL-17 contributes to inflammation, induces reactive astrocytes, and upregulates GFAP and NOS2 (You et al., 2017). We also observed significantly higher expression of IL-22, a strong pro-inflammatory cytokine, which not only upregulates level of inflammatory cytokines but also activates STAT3 transcriptional phosphorylation (Chen et al., 2022). Increase in GFAP and AQP4 in cALD astrocytes along with higher IL-22, IL-17A, and interferon-γ in cALD astrocytes, suggesting that Il-22 in synergy with IL-17 can be a prominent and important mediator of the proinflammatory response in cALD astrocytes.

OSM, a pleiotropic cytokine structurally and functionally related to IL-6, is reported to attenuate inflammatory response and limit tissue damage (Wallace et al., 1995). OSM treatment protected against TNF-α toxicity in experimental autoimmune encephalomyelitis model of multiple sclerosis (Wallace et al., 1999). We observed a synergy is in OSM and IL-6 induction and IL-6 may participate in sustained effects of OSM. OSM is also a late phase cytokine, and this may account for the differential IL-6 levels in AMN astrocytes observed between our and a recent study (Baarine *et al*., 2015). While we measured the OSM/IL-6 levels at 72h the previous study was reported at 6h. Moreover, our expression results are also supported by increased secreted IL-6 protein in the supernatant of AMN astrocytes detected by ELISA. IL-6 is also increased in the plasma of AMN patients (Ruiz et al., 2015). OSM also induces anti-inflammatory gene expression, Arg-1, indirectly through activation of Th2 cytokines in murine macrophages (Dubey et al., 2018). We observed elevated Arg-1 and MRC-1 in AMN astrocytes compared to cALD astrocytes which may relieve inflammatory response. In addition to higher expression of immunoregulatory IL-12α, higher levels of chemotactic CCL2 and CCR2 in AMN astrocytes and increased CXCL1, 2, 8, and 10 may play a beneficial role by inducing recruitment of protective and scavenging microglia to the site of inflammation (Semple et al., 2010).

Inspired by our recent identification of plasma biomarker of disease severity in AMN phenotype (Turk BR, 2022) and to identify upstream regulators of inflammatory pathways, we performed comprehensive miRNA profiling in CTL, AMN and cALD iPSC-derived astrocytes and revealed previously unidentified differentially expressed miRNAs in AMN and cALD astrocytes. Most interestingly, we observed a disease-severity associated increase in miR-9-3p and miR-9-5p expression (CTL<AMN<cALD). miR-9-3p is expressed in response to TLR signaling, and upregulation in activated astrocytes mediates microglial migration and activation via phosphatase and tensin (PTEN) homolog (Yang et al., 2018). miR-9-5p also plays a role in immunometabolic regulation by upregulating cytokine expression and increased glycolysis (Su et al., 2022) similar to our observation in cALD astrocytes. miR-100-5p serves as an extracellular signaling molecule for TLR7/8, and along with miR-298-5p, contributes to neurodegeneration in mouse models of neuronal apoptosis (Wallach, 2021); however, its role in X-ALD is yet to be explored. miR-10a-3p and miR-10a-5p regulate BDNF expression (Zhang et al., 2020a) and the observed higher levels of these miRNA in AMN and cALD astrocytes relate with the lower BDNF expression. IPA analysis predicted IGF1R as target of increased miR-145-5p. IGF-dysfunction is reported in ALD fibroblasts (Al-Essa and Dhaunsi, 2019) and insulin signaling in reduced in *ABCD1*-KO mice spinal cord (Schluter A et al., 2012). Interestingly, we recently documented dysregulated IGF signaling as one of the hubs in plasma metabolite pathways associated with AMN disease severity (Turk BR, 2022). Additionally, we observed upregulated miR-409-3p in AMN and cALD astrocytes. The IPA predicts miR-409-3p targets IL-17A and is regulated by IL-17A, creating a feedback loop. Reports have shown that miR-409-3p contributes to IL-17-induced inflammatory cytokine production in astrocytes via STAT3 signaling pathway in experimental autoimmune encephalomyelitis (EAE) mice (Liu et al., 2019). miR-409-3p expression was also decreased in the plasma of patients with AMN with increasing disease severity (Turk BR, 2022). Similarly, comparative IPA network analysis of CTL vs AMN showed that miR-145-3p is associated with HAND1, ATRX, and RAP2C genes. Altered ATRX and RAP2C have been reported to be associated with gliomas (Alghamri *et al*., 2021; Wang X, 2020). However, further studies are needed to explore the association of miR-145-3p and these genes in X-ALD.

Hematopoietic stem cell transplantation and recently approved gene therapy can arrest cALD disease progression if performed at an early stage (Cartier et al., 2009; Shapiro et al., 2000). However, these therapies cannot recover the function loss already present due to disease progression. Moreover, the mechanism of disease arrest also remains to be explored. We utilized CRISPR/Cas9 knock-in of a functional copy of *ABCD1* in AMN and cALD astrocytes to investigat the changes in the dysregulated molecular, metabolic and miRNA pathways reported above. *ABCD1* induction was sufficient to increase mitochondrial subunits NDUFA6 and SDHB protein levels as seen in OXPHOS immunoblot and SDHB, SDHC, NDUFA6, and UQCRQ in AMN astrocytes suggesting that *ABCD1* expression can partially restore mitochondrial functions. However, cALD phenotype astrocytes did not show restoration in these gene expressions and thus may require additional therapeutic strategies. Interestingly, *ABCD1* knock-in mediated decrease in glycolysis as evidenced by lower expression of key glycolytic genes such as Glut-1, HK-I and -II, GPI, Aldoc, ENO-1, PKM2, and LDHA and a reduced inflammatory cytokine expression in cALD astrocytes suggests that *ABCD1* induction can modulate glycolytic functions. Furthermore, *ABCD1*expression in cALD astrocytes was sufficient to mitigate the basal TLR hyperactivation which is quite intriguing. BDNF and CNTF play essential role in neuronal genesis, differentiation, survival and functional maintenance of sympathetic and parasympathetic neurons (Li et al., 2022; Linnerbauer and Rothhammer, 2020). While AMN-OV astrocytes do not show lowering in glycolysis, BDNF, CNTF, and NGF expressions were increased suggesting that *ABCD1* induction can enhance neuroprotection in AMN phenotype. On the other hand, *ABCD1* induction did not increase BDNF, CNTF, and NGF in cALD astrocytes. Inability of *ABCD1*-overexpression to reverse miR-9-3p and miR-5p expression in cALD astrocytes and thus a sustained inflammation may inhibit these neurotrophic factors in cALD phenotype despite the *ABCD1*-induction.

In summary, cALD astrocytes show increased TLR, MyD88, and NF-κB-expression, which may mediate activation of STAT3, resulting in mitochondrial stress. Our hypothesis is that despite the monogenic mutation in X-ALD, unique inflammatory responses occur in severe cALD and milder AMN phenotypes and involve divergent signaling pathways. A constitutive TLR overactivation may disrupt immune homeostasis, and MyD88-NF-κB-TLR mediated activation of STAT3 then leads to neuroinflammatory and demyelinating X-ALD severity. Activation and induction of ABCD1 expression can mitigate inflammatory burden by lowering the proinflammatory cytokines in AMN and cALD and by lowering TLR expression in cALD astrocytes (Figure 9). Most notably, our study is the first to explore the miRNA landscape of healthy versus diseased X-ALD cells, and our miRNA-seq experiments revealed key differences in post-transcriptional regulatory activity in X-ALD astrocytes. Our findings implicate aberrant STAT3 phosphorylation and elevated miR-9 expression as potential sources of neuroinflammatory responses in cALD, and we observed that OSM may be indicative of the immune landscape in the milder AMN phenotype.

**Figure 9:**
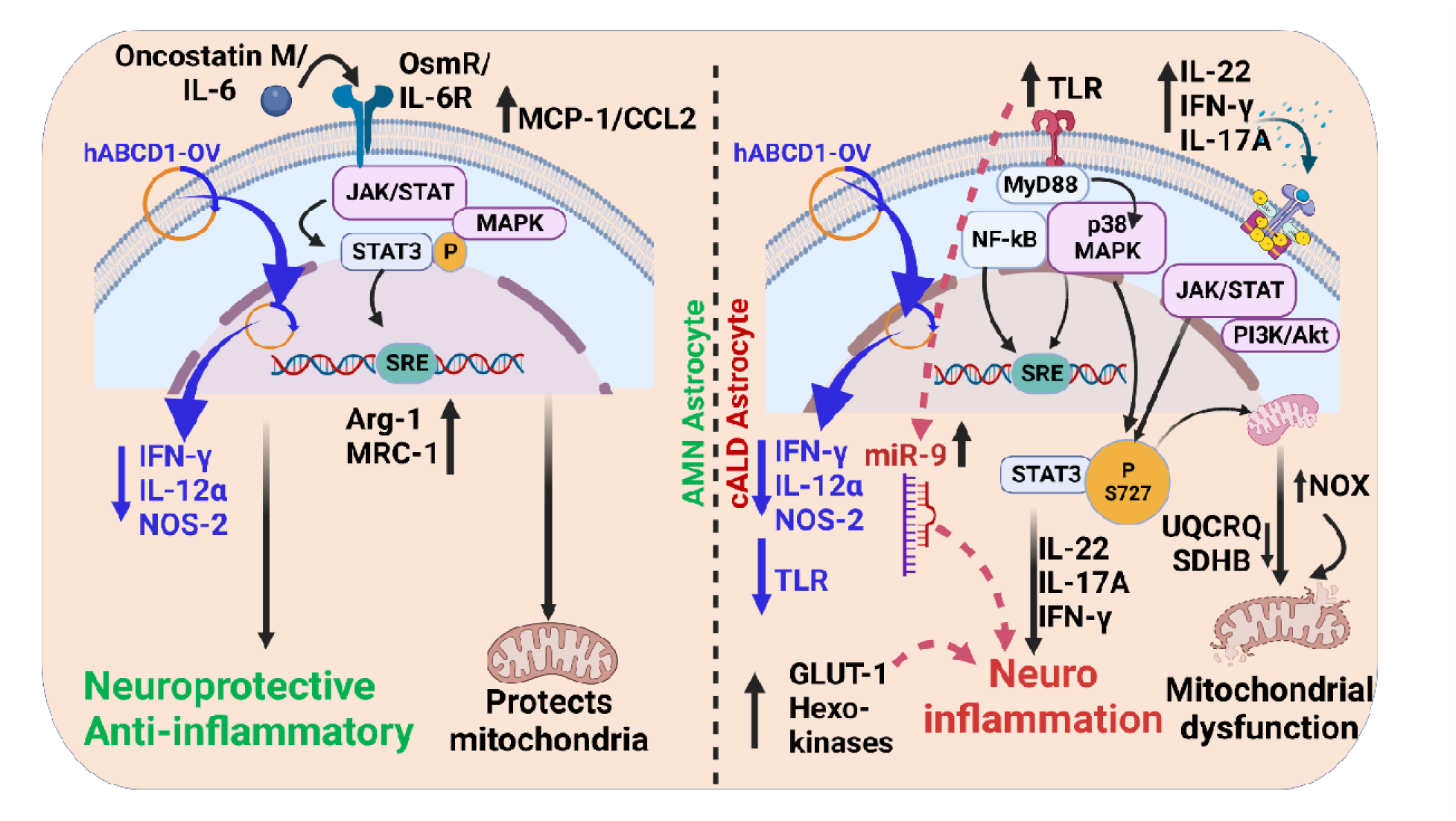
Summarized schematic of study findings and possible mechanisms in AMN and cALD astrocytes. We believe that our findings for the first time show that X-ALD patient-iPSC derived astrocytes can recapitulate molecular and biochemical functional differences and is a robust system to model AMN and cALD phenotypes of X-ALD. In AMN astrocytes, we observed activated Oncostatin-M (IL-6 family) induces Stat3 phosphorylation via gp130-associated JAK/STAT and MAPK and transactivates anti-inflammatory genes especially arginase-1 and mannose receptor C-type-1. Furthermore, higher production of monocyte chemoattractant protein-1 in culture supernatant can be helpful in recruiting protective and scavenging microglia which phagocytose and clear cell debris. On the other hand, cALD patient astrocytes show heightened MyD88-NFκB-TLR activity that fully activate and phosphorylate serine 727 and tyrosine 705-Stat3 resulting in increased proinflammatory cytokines IFN-γ, IL-22, and IL-17A. Elevated NOX expression and reduced SDHB and UQCRQ activity in addition to Stat3 activation is detrimental to mitochondria. Substantial increased TLR activity, mitochondrial dysfunction, and simultaneous hyperglycolytic state are potential contributors to neuroinflammation in cALD astrocytes. miR9 may also be upregulated by activated TLRs and therefore, degrades IL-4 and resulting in reduced anti-inflammatory effect. Furthermore, introduction and induction of *ABCD1* gene using human *ABCD1* overexpression (h*ABCD1*-OV) plasmid in AMN and cALD astrocytes can lower the inflammation. Our findings support our belief that AMN astrocytes show anti-inflammatory (A2) phenotype whereas cALD astrocytes demonstrate highly reactive and proinflammatory (A1) phenotype. **Abbreviations**: AMN: Adrenomyeloneuropathy; Arg-1: Arginase-1; cALD: Cerebral adrenoleukodystrophy; hABCD-OV: Human *ABCD1* overexpression plasmid; MCP-1: Monocyte chemoattractant protein-1; MRC-1: Mannose receptor c-type-1; SRE: Signaling response elements; TLR: Toll-like receptors.

## LIMITATIONS OF THE STUDY

We acknowledge that in light of the variability seen in X-ALD disease, our results for AMN and cALD differential expression warrant validation in iPSCs from larger cohort; however, advanced culture methods have improved the uniformity of accuracy of iPSC cells (Lundin A, 2018). At the time of sampling, the AMN and cALD fibroblasts were identified with respective phenotypes as provided by Coriell Cell repositories. The AMN fibroblast was derived from de-identified 26-year-old patient showing AMN manifested by spastic gait and ataxia, stiff limbs, elevated levels of C26 fatty acids in plasma and fibroblasts. While AMN can progress to cALD up to 30-35 years of age, we do not have information on disease progression of the de-identified patient. Moreover, we and others (Baarine *et al*., 2015; Lee DK, 2019; Ruiz *et al*., 2015; Singh *et al*., 2016; Singh *et al*., 2015) have used the same AMN patient-derived fibroblasts as representative of AMN phenotypes. The lipidomic and transcriptomic analysis of these cells revealed a distinct sphingolipid and triglyceride regulatory factors in AMN not present in cALD (Lee DK, 2019).

## STAR*METHODS

### KEY RESOURCES TABLE

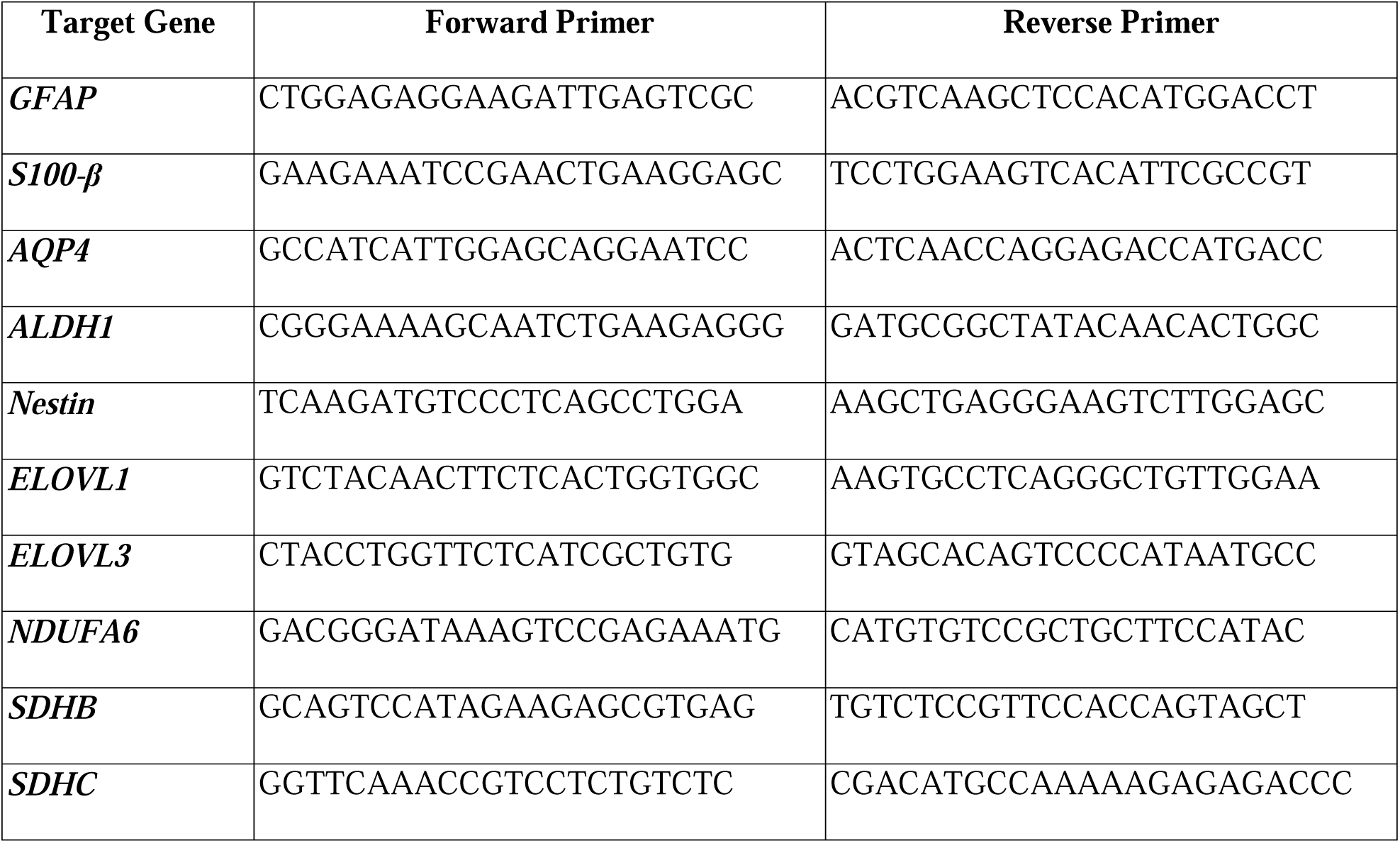
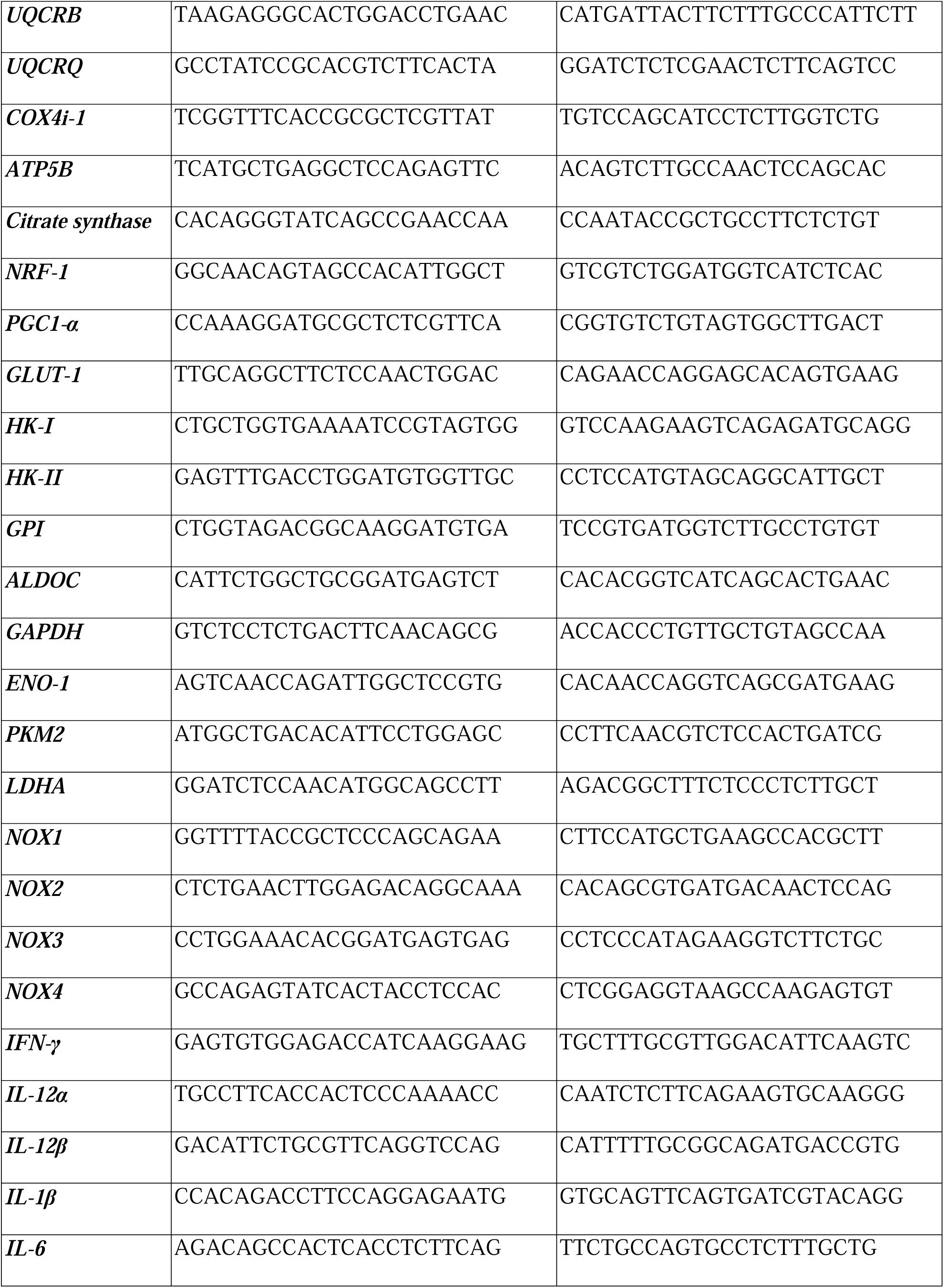
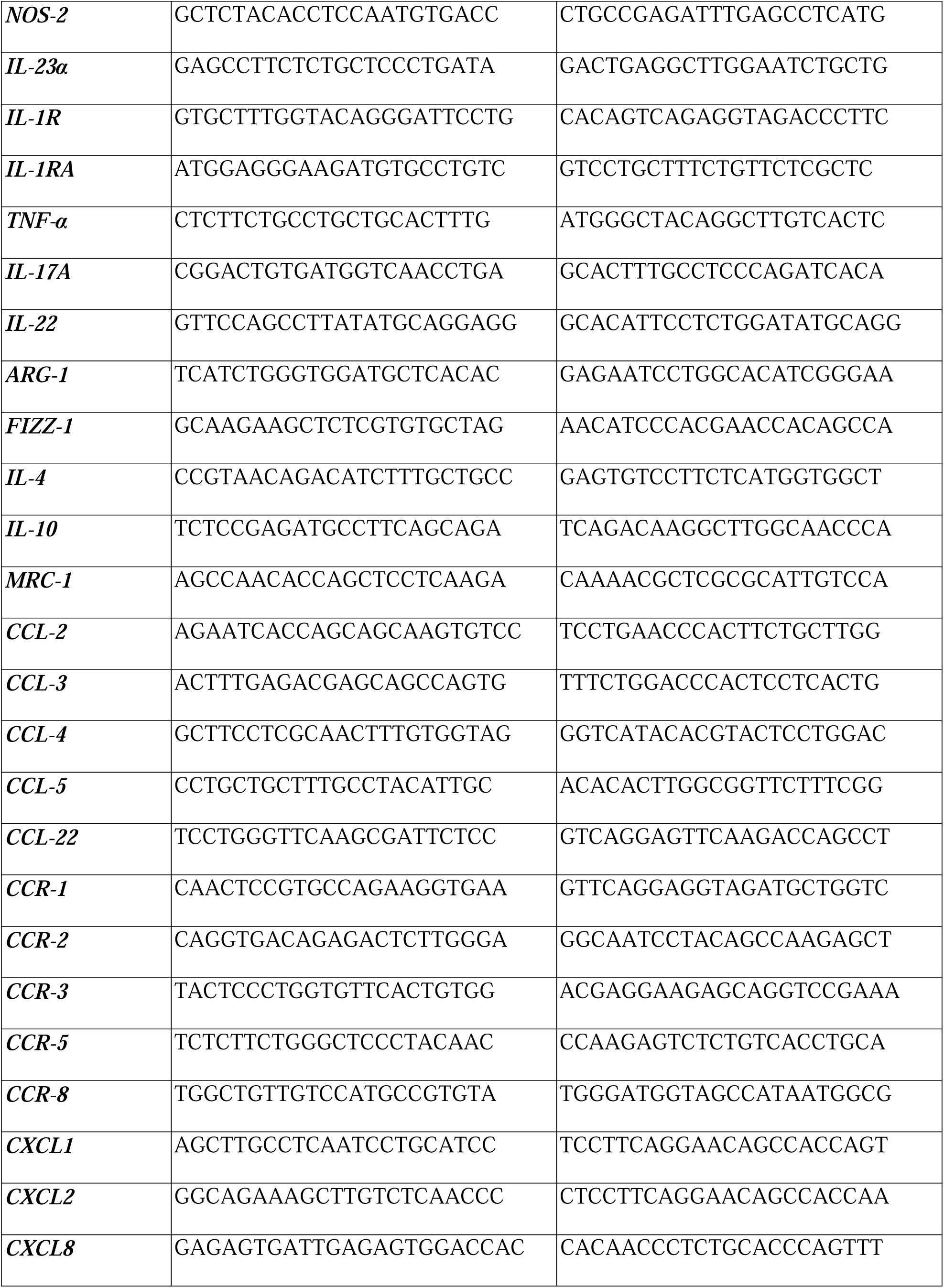
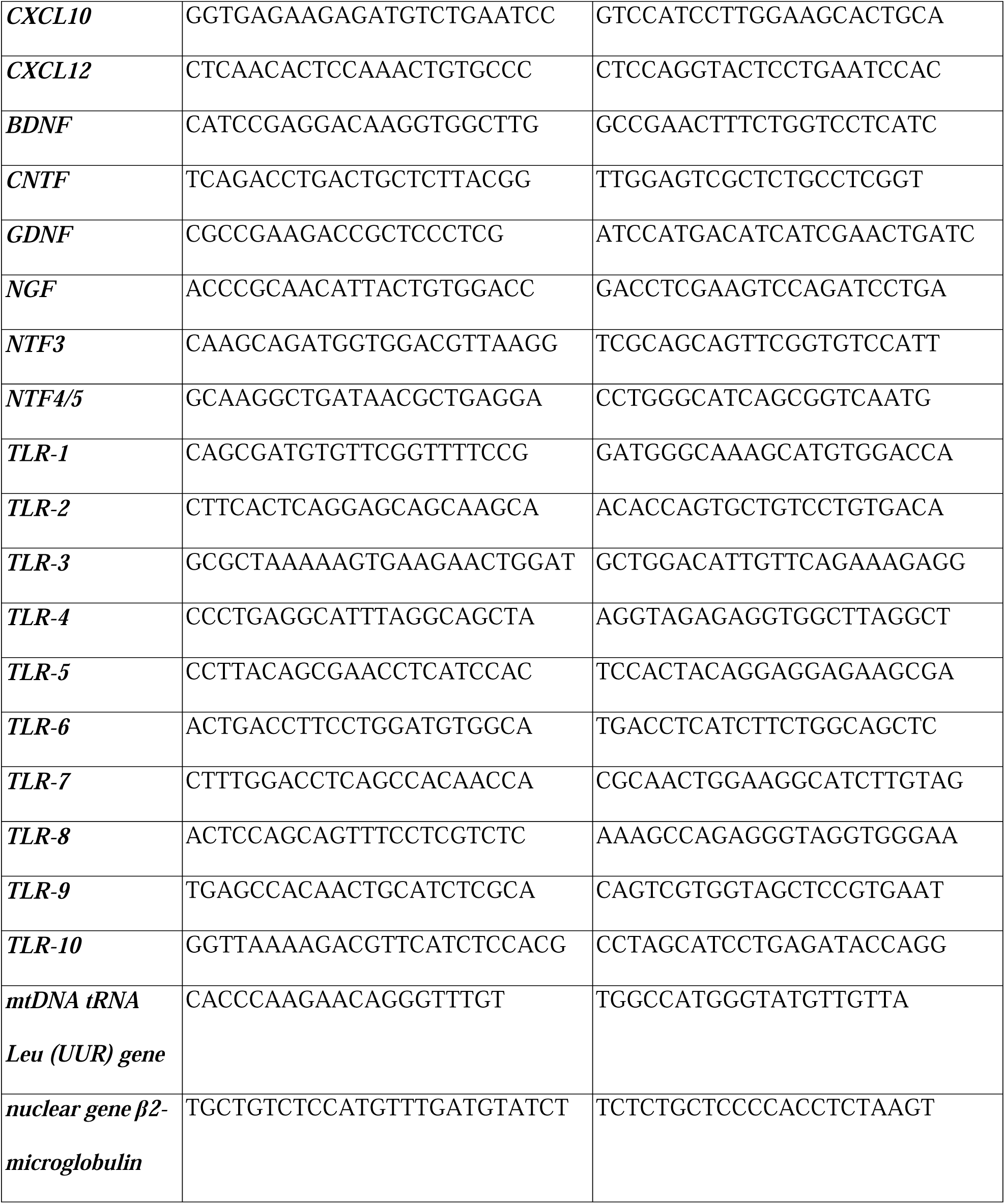
List of primer sequences used for RT-qPCR and Antibodies used for Immunoblotting

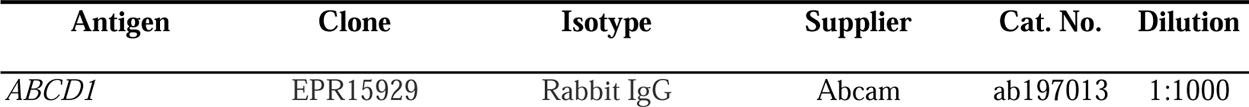
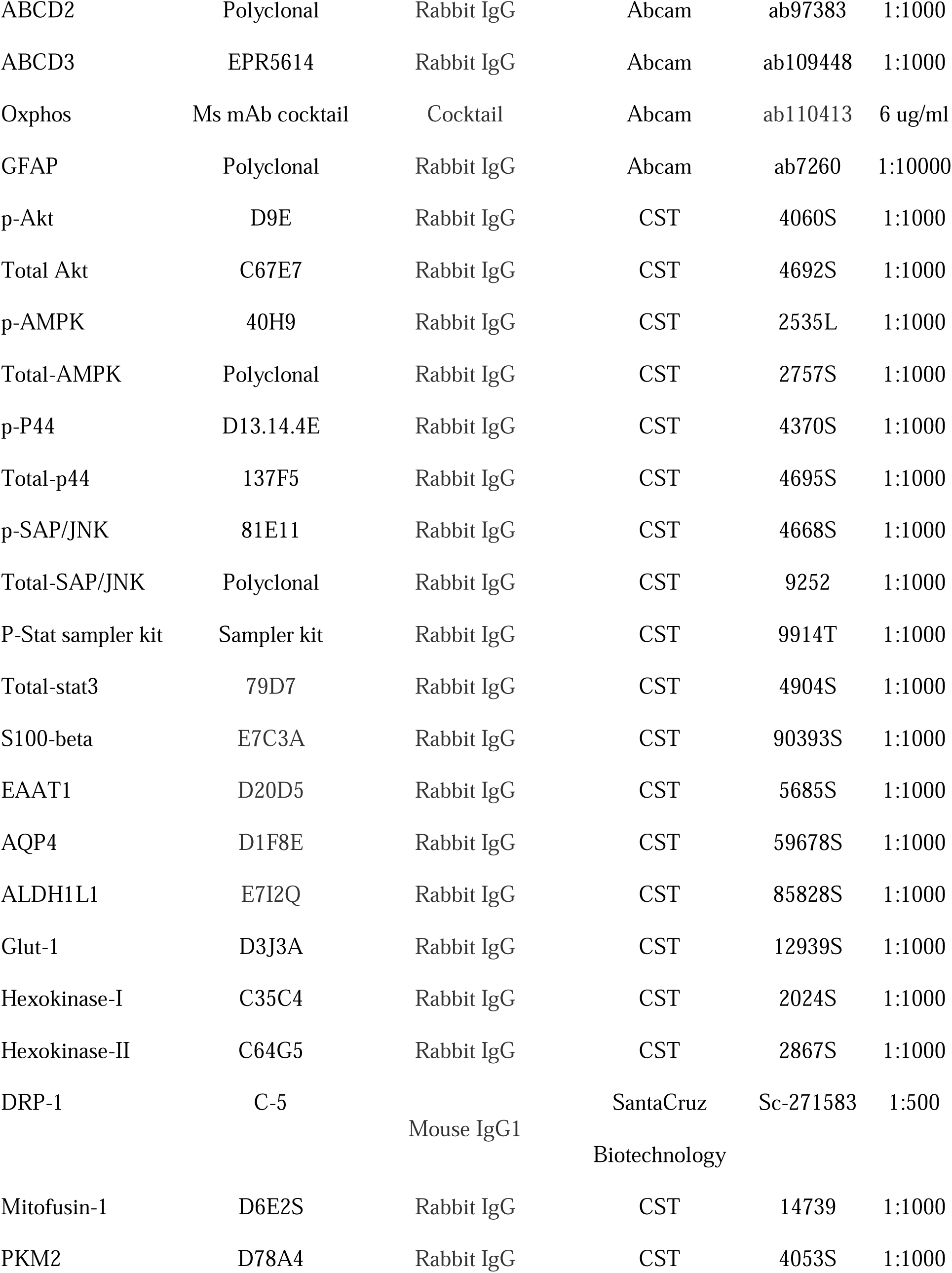
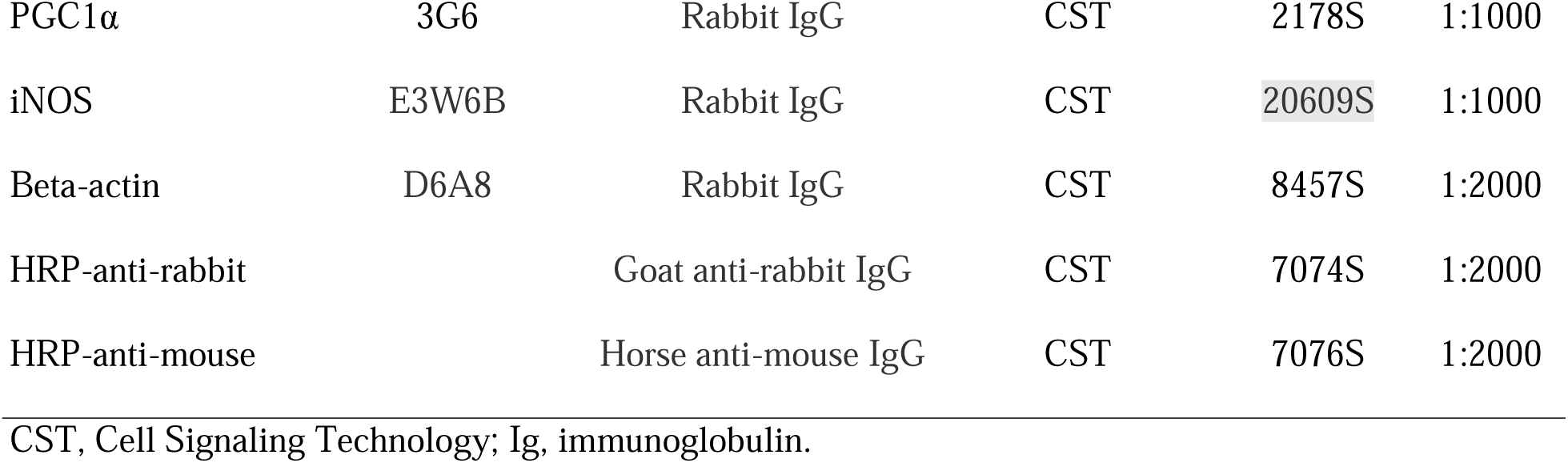

## RESOURCE AVAILABILITY

### Lead contact

Additional information and requests for resources, reagents, and methods should be directed to and will be fulfilled upon reasonable request, by the lead contact, Jaspreet Singh.

### Materials availability

The study did not generate new unique reagents.

## EXPERIMENTAL MODEL AND SUBJECT DETAILS

### Ethics Approval

The project was approved by Henry Ford Health Institutional Review Board (#13352). Fibroblast samples were previously de-identified specimens and did not involve recruitment of human subjects.

### Human fibroblasts

Human skin fibroblasts were obtained from the National Institute for General Medical Sciences human genetic cell repository at Coriell Institute for Medical Research, Camden, NJ. The samples included healthy CTL fibroblasts (GM03377; 19-year-old patient, male), AMN fibroblasts (GM07530; 26-year-old patient, male), and cALD (GM04933; 8-year-old patient, male).

### Derivation of iPSCs and Differentiation into Astrocytes

iPSCs were generated from skin fibroblasts with the Cytotune2.0 Sendai Virus Reprogramming Kit (Axol Biosciences Ltd). Generated iPSCs were differentiated into astrocytes at Axol Biosciences. CTL, AMN, and cALD astrocytes were characterized using immunofluorescence with astrocyte-specific markers. Upon arrival, cells (P5) were frozen in vapor phase nitrogen until used.

## METHODS DETAILS

### iPSC-Derived Astrocyte Culture

Cells were cultured at seeding density of 5 x 10^4^ cells/cm^2^ (day 0) at 37°C and 5% CO_2_ in a humidified incubator in astrocyte medium (Neurobasal-A medium, Thermo Fisher Scientific) containing N-21 max (1X, R&D Systems), fetal bovine serum One Shot (1X, Thermo Fisher Scientific), Glutamax (1X, Gibco, Thermo Fisher Scientific), heregulin-β1 (10 ng/mL, Peprotech, Inc.), basic fibroblast growth factor (8 ng/mL, R&D Systems), penicillin-streptomycin (1X, Thermo Fisher Scientific). For gene expression analysis, cells were cultured in Neurobasal-A medium without serum on Matrigel-coated culture plates. The fibroblasts were characterized and confirmed by nucleoside phosphorylase, glucose-6-phosphate dehydrogenase, and lactate dehydrogenase isoenzyme electrophoresis at National Institute for General Medical Sciences.

### Karyotype Analyses

Human G-banding karyotype analysis was performed using the standard protocol at the Henry Ford Cytogenetics Laboratory. Cells were metaphase-arrested using KaryoMax (Colcemid, Thermo Fisher Scientific) and fixed using a hypotonic (0.8% sodium citrate) and fixative (3:1 methanol:acetic acid) solution. Cells were equilibrated in Thermotron (Thermotron, Inc.) and dried followed by G-banding with Leishman staining. Four metaphases were analyzed per cell type to identify G-banding. Cells were analyzed per the Clinical Cytogenetics Standards and Guidelines(Mikhail et al., 2016).

### Quantitative Real-Time Polymerase Chain Reaction Gene Expression

Total RNA was extracted with the miRNeasy kit (Qiagen) per the manufacturer’s protocol. Single-stranded complementary DNA (cDNA) synthesis and real-time polymerase chain reaction (RT-PCR) were conducted using Bio-Rad’s CFX96 Real-Time PCR Detection System with Bio-Rad IQ SYBR Green Supermix, as described(Singh *et al*., 2009; Singh *et al*., 2016). Primer sets (Integrated DNA Technologies) are listed in Table 1. Gene expression was normalized to 60S ribosomal L27 gene and samples were run in triplicate.

### VLCFA Analysis

CTL, AMN, and cALD astrocytes (2.5 x 10^5^ cells) were submitted to Wayne State University Lipidomics Core facility. Saturated (hexacosanoic [C26:0], tetracosanoic [C24:0]), monounsaturated C26:1, and nervonic C24:1 fatty acids were calculated as ratios against docosanoic (C22:0) fatty acid as internal control. Lipids were subject to alkaline methanolysis and resulting fatty acid methyl esters were analyzed by gas chromatography-mass spectrophotometry (QP2010 GC-MS system, Shimadzu Scientific Instruments) equipped with Restek column, as reported(Singh *et al*., 2016).

### Mitochondrial Oxygen Consumption and Glycolytic Function Measurement

Oxygen consumption rate (OCR) and extracellular acidification rate (ECAR) were measured after 24 hours of culture in 100 mm Matrigel-coated plates (Corning Incorporated) using a Seahorse Bioscience XFe96 Extracellular Flux Analyzer, as reported(Singh and Giri, 2014).

### Western Blot Analysis

Cells were homogenized in radioimmunoprecipitation (RIPA) buffer with Halt protease inhibitor cocktail (Thermo Fisher Scientific). Protein concentration was measured on a Qubit 4.0 fluorometer (Thermo Fisher Scientific) and 50-100 µg of total protein was electrophoresed as described.(Singh *et al*., 2016; Singh *et al*., 2015) Antibodies used are listed in Table 1.

### Transmission Electron Microscopy

One million cells were fixed in 2.5% glutaraldehyde overnight and stained per standard protocol(Zhang et al., 2020b) with slight modifications. Cells were kept in Eppendorf tubes and re-pelleted if needed before infiltration with 1:1 mixture of propylene oxide and Araldite resin (no rocking). Samples were embedded in the mold with resin and cured overnight at 60°C.

Embedded samples were sectioned at 120 nm with an EM U67 Ultratome (Leica Microsystems, Inc.) and collected onto 20 mesh copper grids. Dry grids were stained with uranyl acetate and lead citrate. Images were acquired on a Flash transmission electron microscope equipped with BioSprint camera (Joel USA, Inc.) and image capture software (AMT Imaging Systems).

### Reactive Oxygen Species Production

Reactive oxygen species (ROS) were measured with the Abcam assay kit. Cells (1 x 10^5^) were cultured in triplicate in a 96-well plate and switched to serum free media the next day. Tert-butyl hydroperoxide (50 µM) was added to positive control wells 4 hours before reading. 2,7-dichlorofluorescin diacetate (20 µM) was added to wells and absorbance was read at 485/535 nm.

### Nicotinamide Adenine Dinucleotide Assay

Intracellular nicotinamide adenine dinucleotide/nicotinamide adenine dinucleotide hydrogen (NAD/NADH) assay was performed according to the manufacturer’s protocol (Cayman Chemical). Cells were seeded in 96-well plates (1.0 x 10^5^ cells/well) and incubated overnight at 37°C and 5% CO_2_. Absorbance was measured at 450 nm. NAD/NADH ratio was measured using a NAD^+^ standard curve. Assay buffer served as blank.

### Mitochondrial DNA content

Genomic DNA was isolated using DNeasy Blood & Tissue Kit (Qiagen). Mitochondrial DNA (mtDNA) content was calculated using quantitative RT-PCR by measuring the threshold cycle ratio (ΔCt) of the mitochondria-encoded gene mtDNA tRNA Leu (UUR) gene and the nuclear β2-microglobulin gene (Table 1).(Venegas et al., 2011) Data were expressed as mtDNA/nuclear DNA (nDNA).

### Enzyme-Linked Immunosorbent Assay

Astrocyte serum-free basal culture supernatants were collected after 72 hours, centrifuged to remove residual cells, and stored at –80°C. Enzyme-linked immunosorbent assay Max Deluxe Set (BioLegend) was used to identify interleukin (IL)-6, monocyte chemoattractant protein-1 (MCP-1), C-C motif chemokine ligand 2 (CCL-2), IL-1β, tumor necrosis factor (TNF)-α, IL-12p70, and IL-12/IL-23(p40) following manufacturer’s instructions.

### Assay for Nitric Oxide Synthesis

Nitric oxide production was determined per the manufacturer’s (Sigma Aldrich) instructions using 100 µl lysate in triplicate (assay buffer blanks). Griess reagent I and II were added and incubated for 10 minutes at room temperature. Optical density was measured at 540 nm. Nitrite concentrations were calculated from a standard curve obtained from reaction of nitrogen oxide and Griess reagent II.

### Human Phospho-Kinase Proteome Profiler

We used the human proteome profiler array to detect relative phosphorylation of 37 kinases and 2 related total proteins using 250 µg per the manufacturers’ instructions (R&D Systems).

### Overexpression of *ABCD1* protein

Using CRISPR-Cas9 system, we performed Lipofectamine-mediated (Lipofectamine 2000, Invitrogen) introduction of recombinant *ABCD1* transgene. Briefly, we used 5µg of *ABCD1* overexpression Lenti-viral plasmid (pLV[Exp]-EGFP:T2A: Puro-EF1A> {h*ABCD1*[NM_000033.4, Vector Builder Inc., Chicago, IL) (Figure S2). 5 µg of plasmid was diluted in 150 µl Opti-MEM and Lipofectamine (10 µl) was diluted in 150 µl Opti-MEM media. After 5 min incubation at RT, the diluted plasmid was mixed with diluted Lipofectamine, and the mixture was incubated for 20 min at RT. After 20 min, 250 µl of the complex was added to 1 million cells cultured in media without antibiotics and plated one day prior to transfection. The transgene was checked 48-72 hours after transfection and transfected cells selected with puromycin (0.5ug/mL).

### Data Analysis

Data were analyzed using GraphPad Prism software (version 7.0). Normality was assessed with Kolmogorov-Smirnov test. Groups were compared with two-tailed unpaired Student’s t-test for normally distributed or non-parametric Mann-Whitney test for non-normally distributed data. Statistical significance was set at *P*-value < 0.05.

### Small RNA Library Construction and miRNA-Seq analysis

Small RNA sequencing and informatics was performed by Psomagen, Inc. cDNA library was constructed using QIAseq miRNA Library Kit for Illumina (n = 5). Quality assessment (FastQC v0.11.7), adaptor trimming (Cutadapt 2.8),(MARTIN, 2011) and rRNA filtering were performed. Mature miRNA sequences were mapped to the precursor miRNA sequences of miRBase v22.1 (Bowtie 1.1.2).(Langmead and Salzberg, 2012) miRNA analysis was performed using miRDeep2 (miRDeep2 2.0.0.8)(Friedlander et al., 2012) and R package DEseq2 as published earlier(Turk BR, 2022). Boxplots of miRNA abundance (log10) within the three groups were drawn for altered miRNAs. Partial least squared discriminant analysis was conducted on this subset to visualize the separation between diagnosis groups (R package mixOmics)(Pierre Monget, 2016). The first two components were presented in a scatter plot, with 95% confidence region ellipses plotted for each of the three diagnoses. The common set of altered miRNA was analyzed with QIAGEN Ingenuity Pathway Analysis (IPA; https://digitalinsights.qiagen.com/IPA). Networks of interrelated miRNA and associated molecular components were generated and colored according to observed and predicted levels with QIAGEN IPA.

## DATA AND CODE AVAILABILITY

The data reported in this paper will be shared by the lead contact upon request.

This paper does not report original code.

Any additional information required to reanalyze the data reported in this paper is available from the lad contact upon request.

## ACKNOWLEDGMENTS

The authors recognize the Henry Ford Pathology Surgical Electron Microscopy especially Amy Kemper and Cytogenetics Laboratory especially Dr. Brandon Shaw and Samantha Turpen for their contribution to transmission electron microscopy and karyotyping projects, respectively. We indebtedly thank Dr. Karla Passalacqua, at Henry Ford Medical Writing and Education for her immense help with scientific editing of the manuscript. We also highly acknowledge Stephanie Stebens, at Sladen Library, Henry Ford Hospital for her valuable help with the formatting of manuscript for submission to the journal.

## AUTHOR CONTRIBUTIONS

JS conceived the idea, designed the experiments, provided direction and funding (NINDS grants R21NS114775 and R01NS114245 and Funds from Henry Ford Hospital) for the study. PP, NK, LMP designed and performed the experiments, contributed to the acquisition and analyses of data. PP contributed to drafting the text and preparing the Figure with the help from LMP who helped with miRNA data analysis. All authors revised and approved the final version of the manuscript.

## DECLARATION OF INTEREST

The authors declare that they have no competing interests to disclose.

## FUNDING

This project was supported by National Institutes of Health (NINDS grants R21NS114775 and R01NS114245 to JS and Funds from Henry Ford Hospital to JS). Funding for this project assisted in the ability to collect and analyze our data.

## SUPPLEMENTAL FIGURES

**SUPPLEMENTAL FIGURE S1:**(A & B) RT-qPCR quantification of gene expression for mitochondrial genes (A) and glycolytic genes (B) for CTL, AMN, AMN-OV, cALD, and cALD-OV astrocytes (n = 3 technical replicates per line for astrocytes). (C) RT-qPCR quantification of gene expression for NADPH Oxidases for CTL, AMN, AMN-OV, cALD, and cALD-OV astrocytes (n = 3 technical replicates per line for astrocytes). (D & E) RT-qPCR quantification of gene expression for proinflammatory ((IFN-γ, IL-12α, IL-12β, IL-1β, IL-6, Nos2, IL-23α, IL-1R, IL-RA, TNF-α, IL-17A, and IL-22)) (D) and anti-inflammatory (Arg-1, Fizz-1, IL-4, IL-10, MRC-1, and OSM) (E) in CTL, AMN, AMN-OV, cALD, and cALD-OV astrocytes (n = 3 technical replicates per line for astrocytes). The gene expression of the individual sample was assessed with fold change using the comparative ΔΔ*Ct* method by normalizing with reference gene L-27 in CTL, AMN, and cALD astrocytes. (F, G, & HI) RT-qPCR quantification of gene expression for C-C motif chemokine ligands (CCL-2, CCL-3, CCL-4, CCL-5, and CCL-22), C-C motif chemokine receptors (CCR-1, CCR-2, CCR-3, CCR-5, and CCR-8), and C-X-C motif chemokine ligands (CXCL1, CXCL2, CXCL8, CXCL10, and CXCL12) in CTL, AMN, AMN-OV, cALD, and cALD-OV astrocytes (n = 3 technical replicates per line for astrocytes). (I) RT-qPCR quantification of gene expression for neurotrophic factors (BDNF, CNTF, GDNF, NGF, NTF3, and NTF4/5) for CTL, AMN, AMN-OV, cALD, and cALD-OV astrocytes (n = 3 technical replicates per line for astrocytes). The gene expression of the individual sample was assessed with fold change using the comparative ΔΔ*Ct* method by normalizing with reference gene L-27 in CTL, AMN, and cALD astrocytes. (J) RT-qPCR quantification of gene expression for TLR (TLR-1 thru THR-10) in CTL, AMN, AMN-OV, cALD, and cALD-OV astrocytes (n = 3 technical replicates per line for astrocytes). The gene expression of the individual sample was assessed with fold change using the comparative ΔΔ*Ct* method by normalizing with reference gene L-27 in CTL, AMN, and cALD astrocytes. Color scheme shows colors assigned to sample types in qRT-PCR plots. (L) RT-qPCR quantification of gene expression for hsa-miR-9-3p and hsa-miR-9-5p in CTL, AMN, AMN-OV, cALD, and cALD-OV astrocytes (n = 3 technical replicates per line for astrocytes). The gene expression of the individual sample was assessed with fold change using the comparative ΔΔ*Ct* method by normalizing with reference miRNA RNU6 in CTL, AMN, and cALD astrocytes.

## SUPPLEMENTAL FIGURE S2

Residue: 1-11612 (length: 11612)

Vector Name: pLV[Exp]-EGFP:T2A:Puro-EF1A>h*ABCD1*[NM_000033.4]

Vector Type: Mammalian Gene Expression Lentiviral Vector Vector Size: 11612 bp

Viral Genome Size: 8137 bp Promoter: EF1A

ORF: h*ABCD1*[NM_000033.4]

Marker: EGFP:T2A:Puro

## Notes

### Competing Interest Statement

The authors have declared no competing interest.

### Summary of Updates

X-linked adrenoleukodystrophy (X-ALD) is an inherited progressive metabolic disorder caused by pathogenic variants in the ABCD1 gene, which leads to accumulation of very long chain fatty acids in body fluids and tissues including brain and spinal cord. In the absence of a clear genotype-phenotype correlation the molecular mechanisms of the severe cerebral adrenoleukodystrophy (cALD) and the milder adrenomyeloneuropathy (AMN) phenotypes remain unknown. Given our previous evidence of role of astrocytes in the neuroinflammatory response in X-ALD we investigated the metabolic and molecular profiles of astrocytes derived from induced pluripotent stem cells (iPSC). The iPSCs were in turn generated from skin fibroblasts from healthy controls and patients with AMN or cALD. AMN and cALD astrocytes exhibited lack of ABCD1 and accumulation of very long chain fatty acids, a hallmark of X-ALD disease. Further, cALD astrocytes harbor significantly higher phosphorylation of STAT3, increased Toll-like receptor expression and higher chemokine and cytokine expression. In this first report of miRNA sequencing in X-ALD astrocytes, we observed that miR-9 expression was associated with increasing disease severity phenotype. We performed CRISPR-Cas9 knock-in of ABCD1 gene expression which differentially affected the expression of key molecular, metabolic and microRNA targets in AMN and cALD astrocytes. Extensive characterization of the AMN and cALD iPSC-derived astrocyte model demonstrates critical aspects of X-ALD inflammatory disease in response to ABCD1ABCD1 mutation and can be further utilized for exploring the contribution of astrocytes to differential inflammatory response in cALD.

