## Supplementary figures and images for "iPSC-derived astrocytes to model phenotype-specific differential neuroinflammatory and metabolic responses in X-linked adrenoleukodystrophy"

### FIG S1.tif

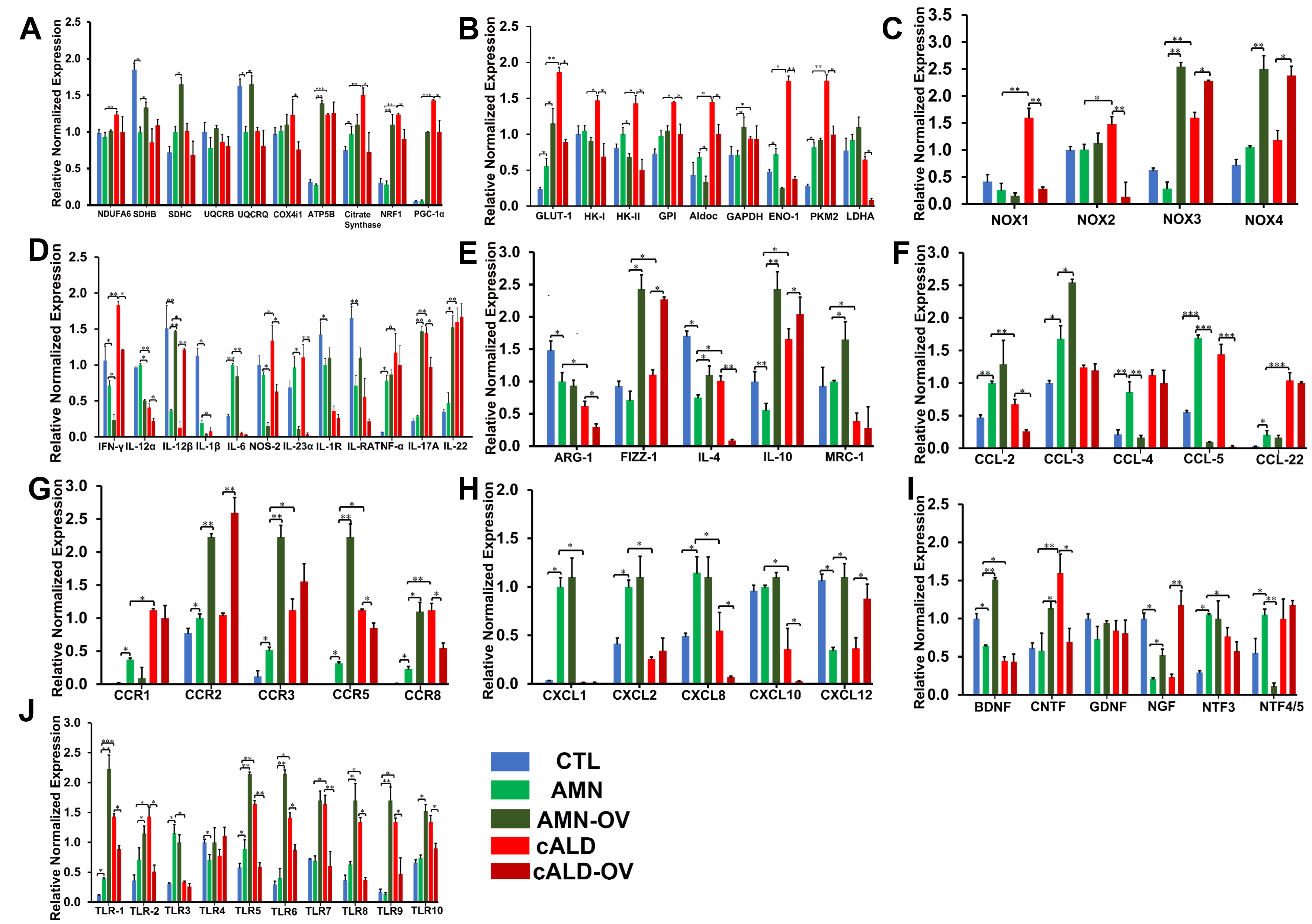

### FIG S2.tif

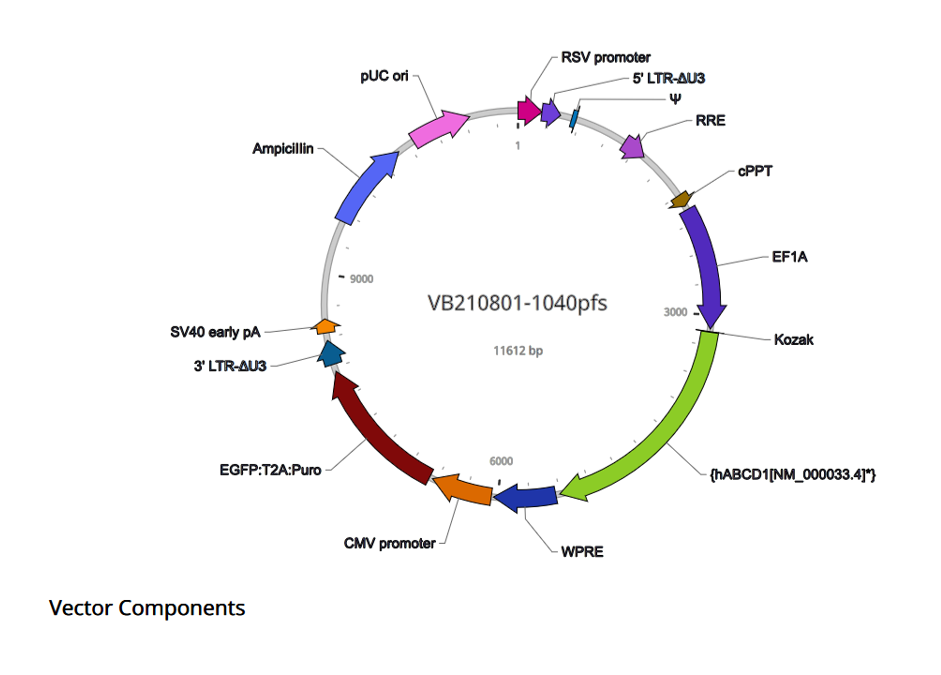
